# The polysaccharide capsule of *Acinetobacter baumannii* affects bacterial adhesion and natural transformation

**DOI:** 10.1101/2024.02.15.580542

**Authors:** Clemence Whiteway, Victor de Pillecyn, Alexandra Maure, Anke Breine, Adam Valcek, Juliette Van Buylaere, Charles Van der Henst

## Abstract

*Acinetobacter baumannii* is an important threat worldwide due to its ability to acquire antibiotic resistance and survive harsh conditions. The polysaccharide capsule represents a major virulence and resistance asset. How the capsular polysaccharides barrier impacts bacterial adhesion remains to be investigated in *A. baumannii.* We characterized capsule mutants of the commonly used AB5075 strain. We assessed how these different capsule mutants adhere to biotic (bacterial and eukaryotic cells) and abiotic surfaces (polystyrene). We confirmed our observations using modern and relevant clinical isolates characterized by different capsule types and capsulation levels. Strains with low capsulation levels systematically depicted increased adhesion compared to capsulated strains, and mucoid isolates showed minimal adhesion. These results show capsule production in *A. baumannii* affects adhesion to various surfaces. We also determined that the presence/absence of *A. baumannii* capsule influences its natural transformability. This illustrates the importance of the trade-off of capsule production in *A. baumannii*.

## Materials and methods

### Bacterial strains and growth conditions

*A. baumannii* strains and plasmids used in this study are listed in Table S1. AB5075-VUB is a clonally isolated strain originating from the parental AB5075 reference strain (1,2). The clinical isolates AB180-VUB, AB183-VUB, AB3-VUB, and AB213-VUB were named according to the field nomenclature by adding the “-VUB” identifier to the strain names and characterized in a previous study (2). Bacteria were grown at 37 °C in 5 mL of liquid broth low salt (LBLS, Luria-Bertani formulation) from Duchefa Biochemie under agitation (166 rcf) or on solid LB-agar plates (1,5%) unless stated otherwise. 30 µg/mL or 50 µg/mL of Apramycin sulfate salt (Sigma–Aldrich) or 10% of crystalized sucrose (Duchefa Biochemie) were used to select/counter select recombinant clones. All experiments were carried out in biological triplicates unless stated otherwise.

### Cloning and generation of capsule mutant of AB5075-VUB

Plasmids and primers used to generate the AB5075-VUB strains used in this study are listed in Tables S1 and S2, respectively. All constructs and plasmids were introduced using electroporation or natural transformation. Primers were purchased from Integrated DNA Technology (IDT), and sequencing was done using the Mix2Seq Kits – Overnight from Eurofins (Sanger sequencing). Gene deletion and complementation were completed following a previously published two-step protocol (3). Gene complementation in the AB5075 chromosome was performed at the predicted neutral Tn7 site (4), and the genes were cloned under the control of the artificial constitutive promoter P_STRONG_ (modified P_tac_ without lacO sites) and the RBS of the super folder GFP (sfGFP). First, the fragment containing the selection/counter-selection cassette *sacB-aaC* cassette flanked by 1–2 kb homologous regions upstream and downstream of the genes of interest (*wza, wzi, or pglL*) was introduced at the targeted locus, and the recombinant clones were selected on apramycin 30 µg/ml. In a second step, the previous clones were transformed with a chimeric product containing the final desired fragment without any marker and counter selected on LB without NaCl (10 g/L tryptone, 5 g/L yeast extract) – agar 10% sucrose and incubated for 6 h at 30 °C. The *wza, wzi, and pglL* coding sequences, the homologous regions flanking these genes, and the *Tn7* site were amplified by PCR from AB5075-VUB genomic DNA. The sfGFP’s coding sequence and P_STRONG_ promoter were amplified from the pASG1 plasmid and the *sacB-aaC* cassette from the pMHL2 plasmid. PCR was performed using PrimeStarMax (TaKaRa), and the resulting products were checked on agarose gel (1%) and purified using the Wizard SV Gel and PCR Clean-Up System (Promega) following the manufacturer’s recommendations. The final chimeric DNA fragments were generated using the Overlap Extension PCR or the NEBuilder HiFi DNA Assembly Master Mix to generate the final chimeric fragments.

### Preparation of Electro-Competent A. baumannii and Electroporation

The clinical isolates AB3-VUB, AB180-VUB, AB183-VUB, and AB213-VUB used in the study are not naturally competent (2). To obtain fluorescent bacteria (GFP) to perform adhesion assays, electro-competent cells from these strains were prepared and subsequentially transformed by electroporation with the pASG1-sfGFP plasmid following an adapted protocol (5). Precultures were started by diluting each strain overnight cultures and incubating at 37°C until an optical density at 600 nm (OD_600_) of 0.4 was reached. All the following steps were performed in the cold. Bacteria were collected by centrifugation for 5 minutes at 5000 rcf; the pellet was resuspended in glycerol 10% and washed twice with 10% glycerol. Finally, the pellet was resuspended in 2 mL of glycerol 10%, aliquoted per 100 µL, and stored at −80°C. Electro-competent *A. baumannii* cells were electroporated in 0.2 mm electroporation cuvettes at 2.5 kV (Ec2 program) with the MicroPuler™ electroporation apparatus (BIO-RAD). Bacteria were cultured in LB for an hour to recover and then plated on apramycin 50 µg/mL.

### Natural transformation of A. baumannii

The AB5075-VUB WT strain and mutants were transformed using an adapted version of the protocol of natural transformation from Valcek *et al*. 2023 (2) with both linear DNA (chimeric DNA to generate the capsule mutants) and the pASG1-sfGFP plasmid. Cultures were initiated from a single clone and were grown at 37°C overnight. The cultures were diluted in PBS to an OD_600_ of ∼0.05. 2 µL of this dilution was mixed with 30-100 ng of DNA (depending on the template) and spotted on 1 mL motility medium (5 g/L tryptone and 2% agar). A negative control without DNA was included for each transformation. The motility media were incubated for 6 h of incubation at 37°C for subsequent selection with apramycin and 6h at 30°C for counter-selection with sucrose 10%. After incubation, 200 µL of PBS was added to each tube and vortexed briefly. 20 and 200 µL of the suspension were plated on a selective medium.

### Transformation frequencies

A hybrid and linear DNA fragment containing (1) an upstream and (2) downstream homologous region with the Tn7 site region of AB5075 and in-between (1) and (2): (3) the *aaC* cassette, encoding for apramycin resistance, was generated by overlap extension PCR to select transformants. The *aac* cassette was amplified from the pASG1-sfGFP plasmid, and the Tn7 site homologous fragments (upstream and downstream) were amplified from AB5075 from genomic DNA using the PrimeStar polymerase (Takara).

Briefly, the AB5075 WT and mutants were transformed using an adapted version of the protocol of natural transformation (Valcek *et al*., 2023) with both the linear *attTn7::aaC* or the pASG1-sfGFP plasmid. Cultures were grown at 37°c overnight in 5 mL of LB. The cultures were diluted in PBS. 30 ng of the *attTn7::aaC* fragment, or 100 ng of the pASG1-sfGFP, were mixed with 10^5^ bacteria in a 2µL volume. This mix was spotted on the motility medium (5 g/L tryptone and 2% agar). A negative control without DNA was included for each transformation. The motility media were incubated for 6 hours at 37°C. After incubation, 200 µL of PBS was added to each tube and vortexed briefly to collect bacteria. The suspension was serial diluted, and 10 µL of each dilution was spotted on both LB-agar with 30 µg/mL of apramycin and on simple LB-agar plates in parallel. The transformation frequencies were obtained by calculating the ratio of transformants (selective media with apramycin) over the total CFUs (LB-agar).

### Galleria mellonella infection

*G. mellonella* larvae were purchased from BioSystems Technology (TruLarv). They were stored at 15 °C and used within the week following arrival. For each strain, 1 mL of overnight culture was washed once in physiological saline (PS: 0.9% NaCl in H_2_O) solution and diluted to a concentration of 1.10^7^ CFU/mL in PS. The larvae were incubated for 30 min at 4 °C before infection to facilitate the procedure. 10 µL of bacterial suspension in PS (1 x 10^5^ bacteria) were injected in the last left proleg of the larvae using 0.3 ml insulin syringes (BD MicroFine). One group of 10 larvae was inoculated per condition. One control group was injected with PS as a negative control, and another group with the virulent AB5075-VUB WT as a positive control of virulence. Survival was monitored for 5 days by checking their melanization and mobility. We stopped monitoring the infection after 5 days because larvae started forming cocoons. Infection experiments were done in biological duplicates.

### Growth measurements

Overnight cultures of each strain were diluted in PBS and plated on solid media (LB-agar) to obtain the colony forming units (CFU) count and determine the correlation between the optical density at 600 nm and the number of bacteria per mL of culture. The growth behavior of the different mutants of AB5075-VUB was assessed in liquid culture using the Cytation 1 plate reader (BioTek, United States). Each well of a 96-well plate was filled with 200 µL of bacterial culture diluted to an initial OD_600_ of 0.1. The OD_600_ was measured every 30 minutes for 24 hours.

### Colony morphology

The opacity of colonies on solid media correlates with the presence or absence of capsular polysaccharides on the bacterial surface of AB5075-VUB (3) by generating respectively an opaque or translucent phenotype. The colony morphology and opacity were assessed by spotting 5 µL of stationary phase bacteria from overnight culture (2-3 x10^9^ CFU/ml) on LB-agar plates (25 ml). After incubation at 37 °C for 24 h, front and back light pictures of the Petri dishes were taken with a Canon hand camera and with the same settings 6 days post inoculation (P.I). Diameters of the macrocolonies were measured using ImageJ. The opacity of colonies at a wavelength of 692 nm was estimated by splitting backlighted pictures of the colonies in a red, green, and blue channel and measuring the mean grey value of each colony in the red channel using ImageJ. The opacity of a colony was calculated by taking the common logarithm of the modal grey value of the media divided by the mean grey value of the colony. This experiment was performed in biological triplicate with technical quadruplicates.

### Density gradient

LUDOX Colloidal Silica (30 wt. % suspension in H2O, Merk) was used to semi-quantify capsule production (3). 1 mL of stationary phase overnight culture was centrifuged for 5 min at 5000 rcf and resuspended in PBS. Then, 750 µL of bacteria in PBS were mixed with 250 µL of LUDOX colloidal silica and centrifuged for 30 min at 9000 rcf. Pictures were taken directly after the centrifugation using a Canon hand camera in front of a black background. The band’s height was measured from the bottom of the tube using ImageJ.

### Pellet stability assay

A simple assay was used to estimate the stability of bacterial pellets after centrifugation and compare this parameter between strains. Briefly, 2 mL of overnight were washed once with PBS, and the OD_600_ was adjusted to 6 for all the strains in a volume of 1 ml. The bacteria were pelleted by centrifugation for 5 min at 5000 rcf. The OD_600_ (OD_i_) of the supernatant was taken as a control to measure the supernatant enrichment after disruption. All the tubes were manually inverted 10 times (upside-down) simultaneously before taking the OD_600_ (OD_f_). The enrichment of the supernatant was determined by calculating the ratio OD_f_/OD_i_. Pictures of the pellets before disruption were taken with a Canon hand camera.

### Culture of cell lines

The A549 human lung carcinoma epithelial cell line (ATCC CCL-185) was cultured in Dulbecco’s Modified Essential Medium (DMEM) supplemented with GlutaMAX™ (Gibco) and 10% heat-inactivated fetal bovine serum (Tico Europe) at 37°C, 5% CO2.

### Adhesion assay to A549 epithelial cells

A549 cells were seeded 18 hours before the experiment in 24-well culture plates at (∼1×10^5^ cells/well, 80% confluence). All the tested strains were transformed with the pASG1-sfGFP plasmid to determine the average number of bacteria (GFP) per epithelial cell (DAPI dye). *A. baumannii* bacteria carrying the pASG1-sfGFP plasmid were grown overnight and centrifuged for 5 min at 5000 rcf and washed 3 times with PBS. Bacteria were diluted in DMEM to infect cells at a <u>m</u>ultiplicity <u>of i</u>nfection (MOI) of 500 bacteria per cell (500:1). Plates were centrifuged at 400 rcf for 4 min and incubated for 1 h at 37°C, 5% CO2. After 1 h, cells were washed 3 times with DMEM and fixed with 4% paraformaldehyde (Thermo Fisher) at room temperature (RT) for 30 minutes. Cells were then stained with DAPI (1 μg/mL, Thermo Fisher) in PBS for 10 min at RT, washed with PBS, and used for acquisition. The MOI was checked by plating the CFU/mL of each inoculum to validate that a similar number of bacteria was added in each condition (**Figure S1**). The number of bacteria per cell was estimated by calculating the total number of bacteria (GFP+ signal) over the total number of cells (DAPI) per picture. ∼500 cells were counted per strain and experiment. Counting was manually done using ImageJ.

### Microscopy

Images were acquired using the Olympus CKX53 inverted microscope (Olympus Life Science) with a 40x objective equipped with a fluorescent illuminator to observe GFP fluorescent bacteria and cell nuclei marked with DAPI.

### Adhesion assay on non-treated polystyrene plates

Overnight cultures were washed once in PBS, and bacterial OD_600_ was measured for all strains tested. Bacterial suspensions were diluted in PBS to OD_600_=0.05 (around 5.10^7^ CFU/mL). 200µL for each strain was added to non-treated 96 well-plates wells (10^7^ bacteria/well). The plate was centrifugated for 2 minutes at 1000 rcf. One picture of the bottom of the well was taken for each strain, as a control that bacteria were in contact with the bottom of the wells and the same proportion. The plate was incubated at RT for 20 minutes. After incubation, the wells were washed 5 times with PBS, and 3 pictures were taken per condition to estimate the number of adherent cells left after extensive washing. The number of cells per picture was estimated using an adapted automated counting method with ImageJ (6). Briefly, the image type was first converted to 8-bit (under “Image” and “Type”) and then to binary-black and white (under “Process” and “Binary”). After that, the threshold (located in the “Image” tab under “Adjust”) was adjusted manually to make sure bacteria were highlighted as black spots (1 spot representing one bacterium equals one particle) with a white background. The parameters were adjusted according to the protocol of Stolze *et al*., 2019 to determine the particle size range and around 15,000 particles/bacteria were counted per picture (under “Analyze”, “Analyze Particles”). The ratio number of bacteria after wash/ before wash was calculated to compare the adhesion of the different strains to polystyrene. The closer values are to “1”, the more adherent the strains are. Statistics were performed on the average fold change value for each tested strain and compared to the average fold change obtained with the AB5075-VUB WT strain.

### Statistical Analyses

All statistical comparisons were performed using GraphPad Prism 9. One-way ANOVA (p = 0.05) test was performed, followed by unpaired t-tests if a difference was observed unless stated otherwise.

The difference is significant if the P-value (P) <0.05. GP(GraphPad): 0.0332<P-value<0.05 (*), 0.0021<p-value<0.0332 (**), 0.0002<p-value<0.0021 (***), p-value<0.0001(****).

## Introduction

*Acinetobacter baumannii* is a Gram-negative bacterium ranked as one of the most concerning bacterial pathogens by the WHO and the CDC, mainly because of carbapenem resistance and its ability to persist and thrive in the clinical setting (Tacconelli et al., 2018; US Department of Health and Human Services & CDC., 2019). While its extensive antibiotic resistance arsenal is well studied, the virulence of *A. baumannii*, mainly based on a “persist and resist” strategy, deserves to be better characterized (Harding et al., 2018). The polysaccharide capsule, among other envelope determinants, is critical to protect the bacteria from the host immune system, desiccation, disinfectants, and antibiotics (Geisinger et al., 2019). Most capsule biosynthesis and export genes are clustered in the capsule locus (K-locus) located on the chromosome. Over 237 K-loci sequences (KL-types) have been identified, illustrating a wide diversity in capsule composition and structure between isolates (Cahill et al., 2022).

While capsule production is regulated, only a few actors have been identified and characterized so far. For example, polysaccharide hyperproduction can be induced during antibiotic exposure through the action of the Two-Component Regulatory Systems (TCS) BfmRS, significantly increasing *A. baumannii* resistance to the complement system and virulence in the murine model (Geisinger & Isberg, 2015). Additionally, phase variation generates phenotypic heterogeneity in genetically homogeneous populations (Ahmad et al., 2019; Tipton et al., 2015). This phenomenon is responsible for the interconversion between opaque, virulent, capsulated bacteria (VIR-O) and translucent, avirulent, and less capsulated (AV-T) bacteria (Chin et al., 2018). The VIR-O variants were more resistant to host-derived antimicrobials and antibiotics. In contrast, the AV-T variants have a decreased virulence *in vivo*, with limited motility but an increased ability to form biofilms (Chin et al., 2018). The specific roles of these two phenotypically different subpopulations in *A. baumannii* virulence and pathogenesis remain to be explored.

Moreover, capsule production can be modulated by Mobile Genetic Elements (MGEs). *A. baumannii*-specific phages often primarily target the capsule, which leads to the emergence of resistant mutants with altered capsule production through IS insertions (Bai et al., 2023; Liu et al., 2022). Indeed, phages have been shown to drive the *in vitro* and *in vivo* evolution of *A. baumannii*, leading to the selection of capsule-deficient but phage-resistant mutants (F. Gordillo Altamirano et al., 2021; F. L. Gordillo Altamirano et al., 2022). In another study, the insertion of ISAba13 into the gtr6 gene induced changes in the capsule structure, leading to increased virulence in mice (Talyansky et al., 2021). Our recent study characterized a natural non-capsulated mutant with an ISAba13 element interrupting the itrA gene. We showed that upon stress exposure with antibiotics, scarless excision of ISAba13 restored capsule production and virulence (Whiteway et al., 2021).

The capsule is a critical virulence and resistance factor for several bacterial pathogens, including *A. baumannii*. However, polysaccharide capsules have been shown to impede adhesion by shielding the functions of adhesins and fimbriae on the bacterial surface, preventing invasion of host cells and biofilm formation (Béchon et al., 2020) of several pathogenic bacteria such as *Klebsiella pneumoniae* (Schembri et al., 2005), *Neisseria meningitidis* (Spinosa et al., 2007), *Haemophilus influenzae* (Häuser et al., 2018) and *Streptococcus pneumoniae* (Hammerschmidt et al., 2005). Adhesion is the critical first step for interactions with biotic and abiotic surfaces, conditioning successful invasion of new environments and survival of numerous commensal and pathogenic bacteria. This complex process depends on multiple physical and chemical factors from the surface, bacterial cells, and their physiology (Ellison, 2018). However, how *A. baumannii* capsule production impacts the adhesion to the biotic and abiotic surfaces remains unclear and has never been investigated *per se*. The same is true for the influence of *A. baumannii*’s capsule on its natural transformation. It was shown that capsules could limit processes such as natural competence with different bacterial models (Jeon *et al*., 2009; Marks *et al*., 2012; Schaffner *et al*., 2014). However, in a recent bioinformatic study, analysis of thousands of bacterial genomes revealed a positive correlation between the presence of a capsule locus and the frequency of genetic exchanges by diverse means (Rendueles *et al*., 2018). Mechanisms explaining this correlation were not identified but this result suggests that, in some cases, a mechanism directly linked to capsule production could increase genetic exchanges.

Our study focuses on understanding how the presence or absence of the polysaccharide capsule impacts the adhesion of *A. baumannii* to biotic and abiotic surfaces and its natural transformation ability. We generated different capsule mutants of the AB5075-VUB (KL25) strain and characterized their capsule production and virulence phenotypes *in vivo*. We also used four modern clinical isolates characterized in a previous study with different capsulation levels and capsule types (KL): AB3-VUB (KL40), AB180-VUB (KL3), AB183-VUB (KL9), and AB213-VUB (KL40) (Valcek et al., 2023). The AB3-VUB and AB213-VUB strains were constitutively mucoid, and AB180-VUB and AB183-VUB display limited capsulation. We assessed their ability to form cohesive pellets and to adhere to polystyrene plates and pulmonary epithelial cells. This project aims to understand better the polysaccharide capsule’s impact on the adhesion process in *A. baumannii*.

## Results

### Phenotypical characterization of AB5075-VUB capsule mutants and clinical isolates of *A. baumannii*

The commonly used AB5075 strain belongs to the serotype KL25, producing a substantial capsule that can be over-produced during antibiotic treatment (7). To assess the impact of the presence/absence of a capsule, we generated deletion strains of the *wza* and *wzi* genes from the late stage of the Wzy-dependent pathway (8). Wza is an outer membrane protein essential for the export of capsular polysaccharides (CPS) (9). Wzi is required to strongly associate CPS to the bacterial cell surface (10). We included the deletion strain of *itrA* already published in a previous study (3). ItrA is the initial glycosyltransferase required to construct both the glycan repeat units that form the capsular exopolysaccharide and O-linked protein glycans (adding the first sugar to the UndP carrier) (11). Therefore, we additionally generated a deletion strain of *pglL* as a control to exclude the fact that the phenotypes observed with the Δ*itrA* strain are linked to the O-linked protein glycosylation rather than capsule formation. We generated complementation strains for all the deletions mentioned above by cloning the respective gene at the predicted neutral Tn7 site (4) of the AB5075 chromosome under the control of a constitutive promoter (see Materials and methods).

We semi-quantified capsule formation using a silica colloidal density gradient previously validated (3). Briefly, capsulated bacteria form bands in the higher fraction of the gradient after centrifugation due to their lower density, whereas non-capsulated ones will be in lower fractions because of their heavier densities. We took pictures of the density gradients (Figure 1A) and determined the average height of the bands from the bottom of tubes (Figure S1) for all the strains used in this study. Deletion of *wza* or *itrA* abolishes capsule production, while deletion of *wzi* leads to an intermediate phenotype. Deletion of *pglL* does not affect capsule production in the tested conditions. The respective complementations at the Tn7 site restore capsule production to the WT level. We detected a slightly but statistically significant higher band for the Δ*wzi*/*wzi* strain that could be an effect of the constitutive promoter used for gene complementation (see Material and Methods).

**Figure 1.**
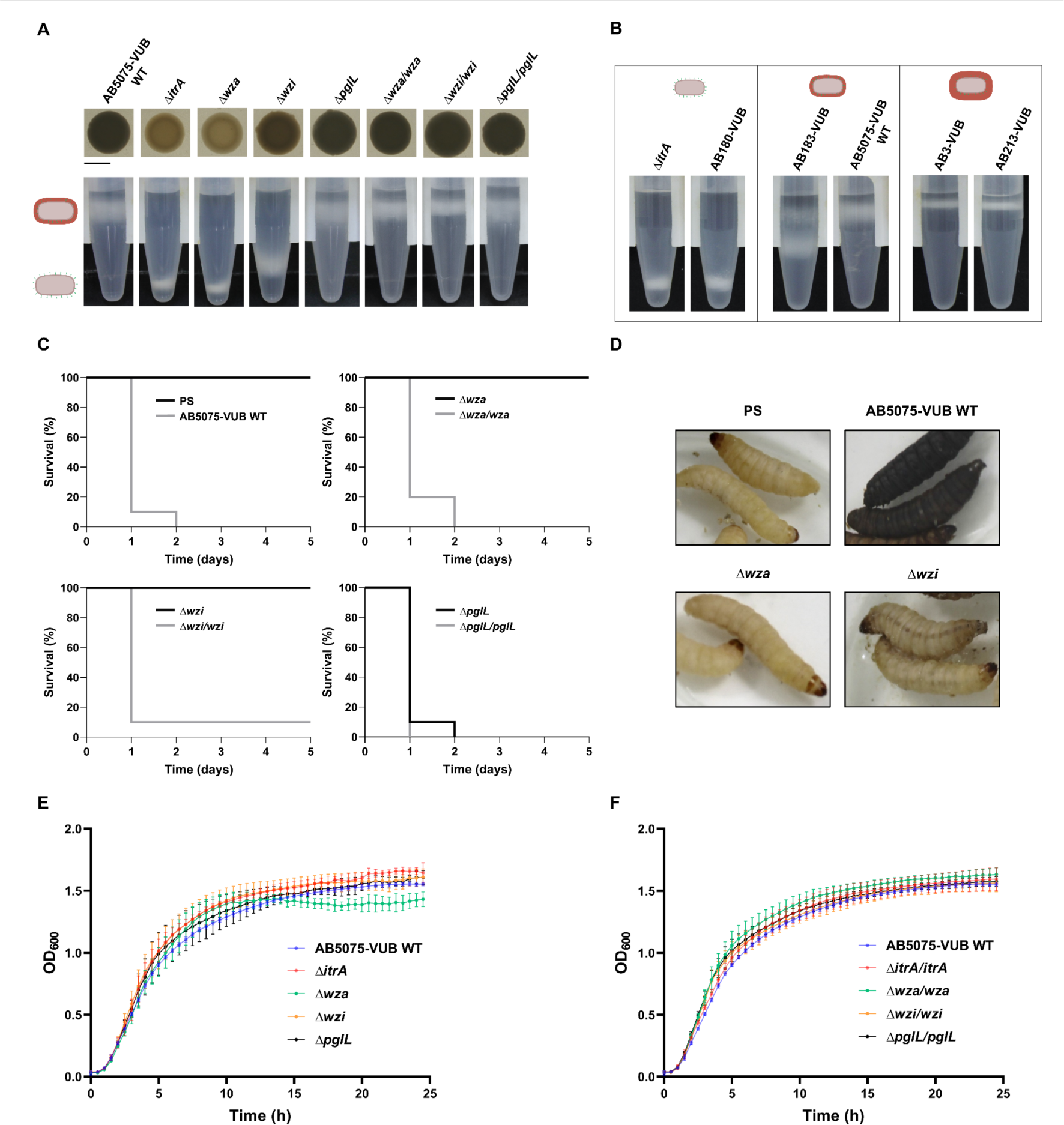
Phenotypical characterization of AB5075-VUB capsule mutant and clinical isolates of *A. baumannii:* (A) Macrocolony opacities and density gradients of the AB5075-VUB capsule mutants. Scale bar: 0.5 cm. (B) Density gradients of the clinical isolates of *A. baumannii*. The strains are categorized into 3 groups: low capsulation, intermediate capsulation, and high capsulation (here constitutively mucoid), with respective cartoons depicting the capsulation level of bacterial cells. (C) *In vivo* virulence of AB5075-VUB capsule mutants in *G. mellonella*. Survival of the larvae was followed over 5 days: the y-axis corresponds to the percentage of larvae survival (%), and the x-axis corresponds to the time post-inoculation (days). Ten larvae were infected for each condition. (D) Pictures of *G. mellonella* larvae showing their phenotype one day post-infection with AB5075-VUB WT and capsule mutants. Darkening of the larvae corresponds to complete or partial melanization. (E) Growth measurements of AB5075-VUB WT and deletion mutants. OD_600_ measurements over time for 24h; (F) Growth measurements of AB5075-VUB WT and complementation strains. OD_600_ measurements over time for 24h. Error bars indicate the standard deviation (s.d) for the three biological replicates of each strain.

We also included four additional clinical isolates from our collection (12) that are already characterized, including for capsule production and virulence in *Galleria mellonella* (2). Strains used in this study present different KL types determined according to their K-locus genetic content and structure (13). The capsule type associated with the AB180-VUB strain is KL3, KL9 for AB183-VUB, KL40 for AB3-VUB and AB213-VUB (2). The WT AB5075-VUB strain (KL25 type) is a positive control for capsule production and Δ*itrA* for the absence of a capsule. We used an adapted protocol of the density gradients from a previous study (see Material and methods) (3) by adapting the Ludox concentration, generating new capsulation-level categories for each strain (Figure 1B). We measured the average height in the gradient for all the isolates (Figure S1B). AB180-VUB (KL3) displays low capsulation, and we see a similar band as with the non-capsulated ΔitrA in the gradient, confirming former observations (2). For AB183-VUB (KL9), TEM pictures showed a low capsulation as well (2). However, this gradient’s result is less categoric (Fig. 1B), and we observe two distinct bands using this new Ludox concentration. The AB183-VUB phenotype can be classified as intermediate capsulation since the band is in the middle part of the tube and is quite dispersed. This distribution could be the hallmark of a heterogeneous production of capsules in the AB183-VUB strain. As previously described with TEM pictures and density gradients, AB3-VUB (KL40) and AB213-VUB (KL40) strains produce a large capsule that is additionally witnessed by a constructive mucoid phenotype on solid media (2,14) and will be classified as highly capsulated (high and thin band in the gradient).

Capsule production correlates with macrocolony opacity of the AB5075-VUB WT strain and associated mutants on solid media (Figure 1A and Figure S2A). As previously reported, non-capsulated variants (here Δ*itrA* and Δ*wza*) of AB5075 show a translucent phenotype, whereas capsulated ones show an opaque phenotype (3). As expected, we observed an intermediate opacity for the Δ*wzi* strain, correlating with the phenotype in the density gradient (Figure 1A and Figure S2B). Upon deletion of *pglL* and complementation of all the deleted genes, all strains display an opacity level similar to the WT phenotype. We determined the opacity of the macrocolonies six days post-inoculation (P.I) on solid media. The Δ*wzi* shows a decreased opacity when all the other strains have a similar or increased opacity on day 6 compared to day 1 (Figure S2C and D). This could indicate a loss of capsule between days 1 and 6. This result supports that the Wzi outer membrane protein is essential for capsule stability over time (10).

Next, we assessed the *in vivo* virulence of the different mutants and respective complementation strains using *Galleria mellonella* larvae (Figure 1C). Deletions of *wza* and *wzi* impair virulence, and larvae survive after 5 days following inoculation. Though most of the larvae infected with the Δ*wzi* mutant survive after 5 days, we observed signs of toxicity through partial melanization across the tail/headline and reduced mobility of the larvae (Figure 1D) already one day after infection. This phenotype was not observed in the inoculation of Δ*wza* bacteria or with the negative control (PS). Deletion of *pglL* does not impact the virulence in the tested conditions. All the complementation strains show virulence like the WT strain, with most larvae dying within one to two days post-infection.

We monitored the growth of the different mutants in a rich liquid medium (LB) over 24 hours to complete their characterization and test for potential growth defects. All AB5075-VUB deletion and complementation strains (Figure 1E and F) grow similarly to the WT, except for the Δ*wza* mutant, for which we observed a growth defect in the stationary phase (Figure 1E). However, we did not detect a significant difference in the colony-forming units generated overnight (Figure S3). Since we confirmed that mutations in the late stage of capsule biosynthesis can lead to toxicity, we decided to systematically use the Δ*itrA* and Δ*pglL* mutants in parallel with the Δ*wza* mutant (15).

### Inter-bacterial cohesion forces are stronger in the absence of capsule

During routine experiments, we observed that the strain AB5075-VUB WT pellet is consistently larger than the pellet observed with the non-capsulated Δ*itrA* mutant after centrifugation, despite using an equal number of bacteria (Figure 2A). We hypothesized that the capsule may be involved in this phenotype. Hence, we compared the area of the pellets generated by the different mutants (Figure 2A and C) and the clinical isolates (Figure 2B and D). The non-capsulated strains with low capsulation have smaller pellets than capsulated ones, and the mucoid strains AB3-VUB and AB213-VUB significantly display the biggest pellets (Figure 2B). Therefore, the size of the pellets in the tested conditions positively correlates with the capsule production level (Figure 2C and D)

**Figure 2.**
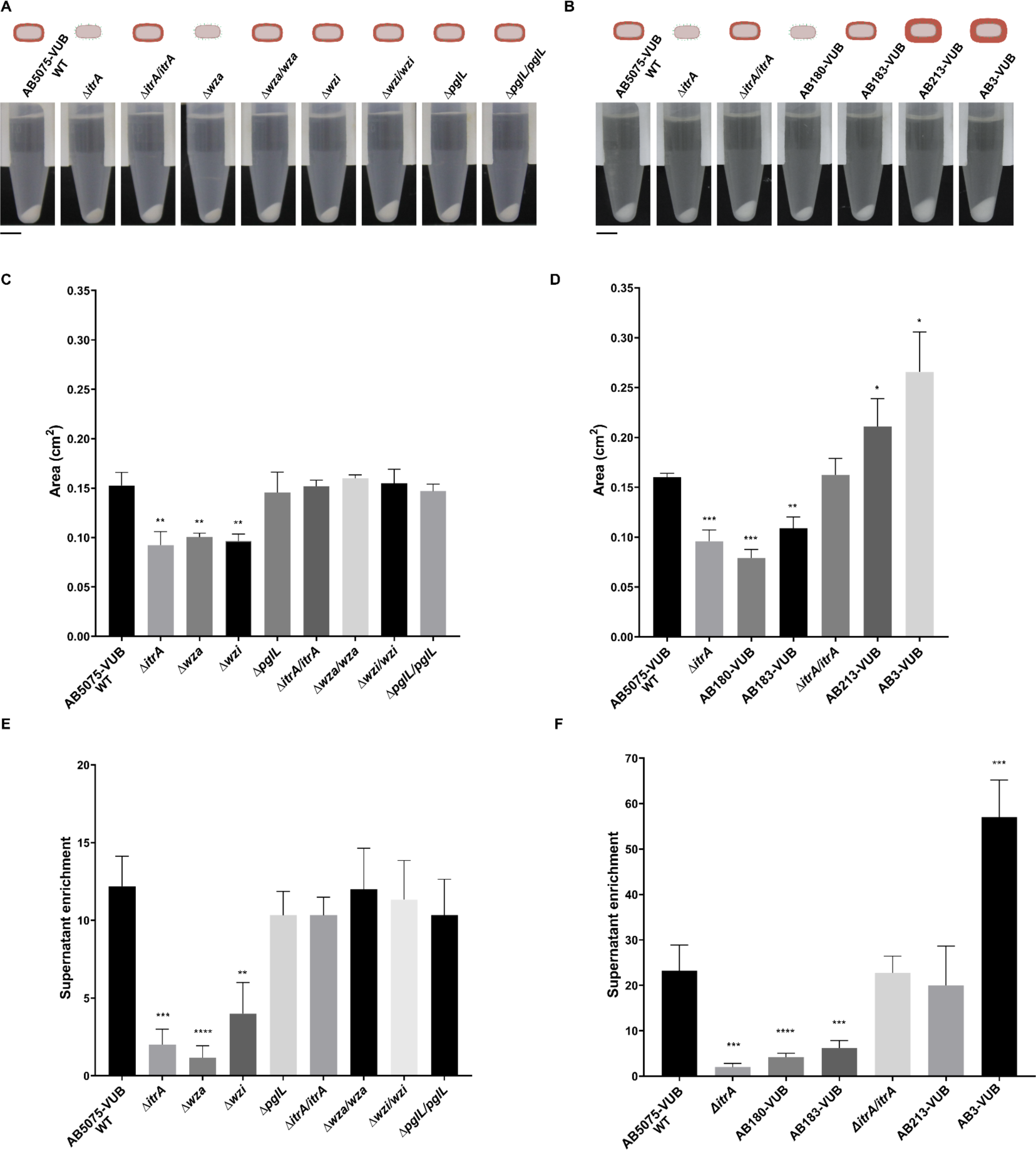
Cohesion forces between *A. baumannii* bacteria in pellets. (A) Pellets pictures of the AB5075-VUB capsule mutants and (B) clinical isolates after centrifugation. Scale bar: 0.5 cm. (C) Area of the pellets measured for the AB5075-VUB mutants (cm^2^) and (D) for the clinical isolates (cm^2^). (E) Supernatant enrichment of the AB5075-VUB capsule mutants and (F) the clinical isolates. A one-way ANOVA was performed for each set of experiments and showed significant differences among the means for the supernatant enrichment. Then unpaired student tests (t-tests) were performed on the data by comparing each strain to the AB5075-VUB WT (reference). The difference is significant if the P-value (P) <0.05. GP(GraphPad): 0.0332<P-value<0.05 (*), 0.0021<p-value<0.0332 (**), 0.0002<p-value<0.0021 (***), p-value<0.0001(****).

As the pellets contain the same number of bacteria, we hypothesized that smaller pellets might reflect higher interaction forces between bacterial cells. To test that hypothesis, we centrifugated the bacteria and measured the enrichment of the supernatant in bacteria by flipping the tubes upside down 10 times (see Material and Methods). We measured the optical density of the supernatant before (ODi) and after (ODf) homogenization. The ratio ODf/ODi, representing the enrichment of supernatant after homogenization is presented in Figure 2E for the AB5075-VUB mutants and Figure. 2F for the clinical isolates. We observed a high and similar enrichment for AB5075-VUB WT, Δ*pglL*, and all the complementation strains. However, we see a low enrichment of the supernatant after flipping with the Δ*itrA* and Δ*wza* mutants (respectively, 6-fold and 10-fold decrease enrichment compared to the WT) and an intermediate enrichment for the Δ*wzi* mutant (3-fold decrease compared to the WT). The constitutively AB3-VUB displays a highly unstable pellet (2-fold increase in the enrichment compared to AB5075-VUB WT), whereas the other mucoid isolate AB213-VUB depicts a similar enrichment as AB5075-VUB WT after flipping. As expected, the supernatant enrichment is low for AB180-VUB and AB183-VUB with limited capsulation (6 and 4-fold decreases, respectively, compared to the AB5075-VUB WT). These observations show that pellets formed by capsulated bacteria are much less stable than those of non-capsulated ones, pointing toward different levels of bacterial adhesion forces.

### The capsule of *A. baumannii* impairs adhesion to eukaryotic cells

Bacterial adhesion is the first and critical step for surface colonization and is consequently crucial for cell invasion and pathogenesis. Consequently, we assessed how the different capsule mutants and the clinical isolates of *A. baumannii* adhere to pulmonary A549 epithelial cells. Prior experiments confirmed that the carriage of the pASG1-sfGFP plasmid does not impact the growth of AB5075-VUB and that 99.6% of the cells transformed were expressing green fluorescence (Figure S4A and B). CFUs were determined for all the inoculums to make sure that a similar number of bacteria was added to each well with all the strains (Figure S5). The non-capsulated strains are more adherent than the capsulated strains in the tested conditions (Figure 3A and B). Indeed, the Δ*itrA*, Δ*wza* strains are significantly more adherent than the WT, with respectively a 36-fold and 53-fold increase in adherence. Δ*wzi* showed an intermediate phenotype, with an average 15-fold increased adherence compared to the WT. This is consistent with the intermediate phenotype shown previously (Figure 1A). Deletion of *pglL* has no detectable effect on the adhesion in the tested conditions. and all the complementation strains display an adhesion level similar to the WT.

**Figure 3.**
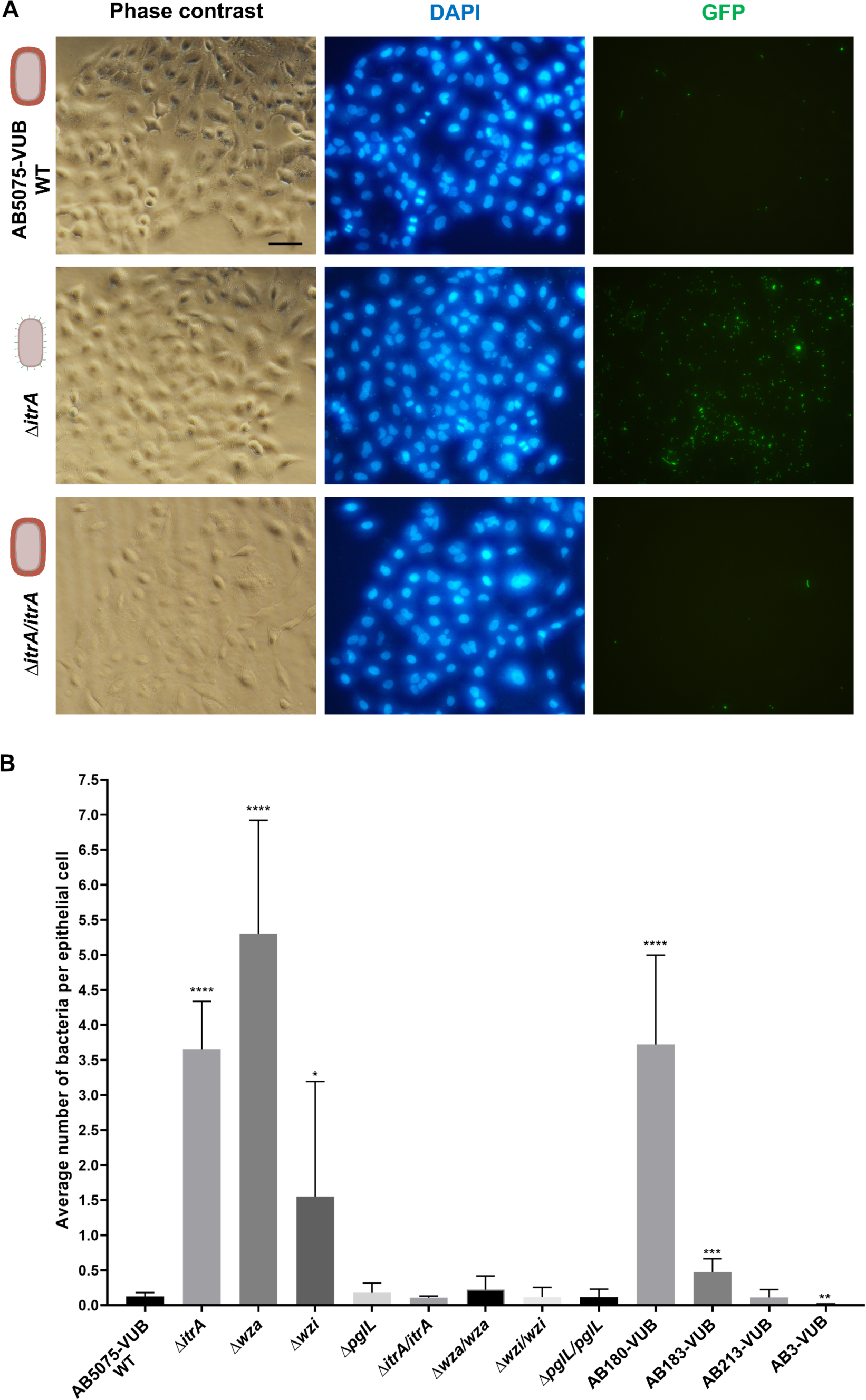
Adhesion of *A. baumannii* to epithelial cells. (A) Representative microscopy pictures of the adhesion assay for AB5075-VUB WT, *ΔitrA,* and *ΔitrA/itrA.* Scale bar: 50 µm. (B) Quantification of the adhesion of different capsule mutants and clinical isolates to A549 epithelial cells represents the average number of bacteria per epithelial cell for each strain. A one-way ANOVA was performed for each set of experiments and showed significant differences among means (number of bacteria per epithelial cell). Then unpaired student tests (t-tests) were performed on the data by comparing each strain to the AB5075-VUB WT (reference). The difference is significant if the P-value (P) <0.05. GP(GraphPad): 0.0332<P-value<0.05 (*), 0.0021<p-value<0.0332 (**), 0.0002<p-value<0.0021 (***), p-value<0.0001(****).

Concerning the clinical isolates, the low-capsulated AB180-VUB, like the non-capsulated mutants Δ*itrA* and Δ*wza*, adheres better compared to AB5075-VUB WT with a 37-fold increase in adhesion (Figure 3B). AB183-VUB shows a limited adhesion but was significantly higher than the one observed with AB5075-VUB WT (4-fold increase), which is consistent with the TEM pictures of the labelled capsule from our previous work, with limited capsulation level (2). AB213-VUB shows a similar adhesion as AB5075-VUB WT, while AB3-VUB has very limited adhesion abilities (a 10-fold decrease). Microscopy pictures of the adhesion assay for the capsule mutant and clinical isolates are provided in Figure S6. Taken together, these results indicate that bacterial adhesion to eukaryotic cells is impaired by capsule production in *A. baumannii*, in the tested conditions.

### The mucoid phenotype inhibits adhesion to abiotic surface

We then assessed the adhesion of all the strains on abiotic plastic surfaces (Figure 4A). First, we observed that the AB5075-VUB WT strain is intrinsically adherent to polystyrene in the conditions used in this experiment (Figures 4A and 4B). However, when we calculated the ratio of cells after versus before wash (Material and Methods), we observed that *ΔitrA*, *Δwza,* and *Δwzi,* are more adherent compared to the AB5075-VUB WT and the complementation strains. We consistently observe variable patterns (homogenous or clustered deposition) in the wells for capsulated AB5075-VUB WT, *ΔpglL,* and the complementation strains (Figure 4A and S7). The low-capsulated AB180-VUB is highly adherent in the tested conditions, and the constitutively mucoid strains AB3-VUB and AB213-VUB show very low levels of adhesion (Figure 4A, 4B, and S7). We observed an intermediate phenotype for the AB183-VUB strain, similar to the AB5075-VUB capsulated strains. Hyper-capsulation, leading to a mucoid phenotype, almost abolishes adhesion to polystyrene in the tested conditions.

**Figure 4.**
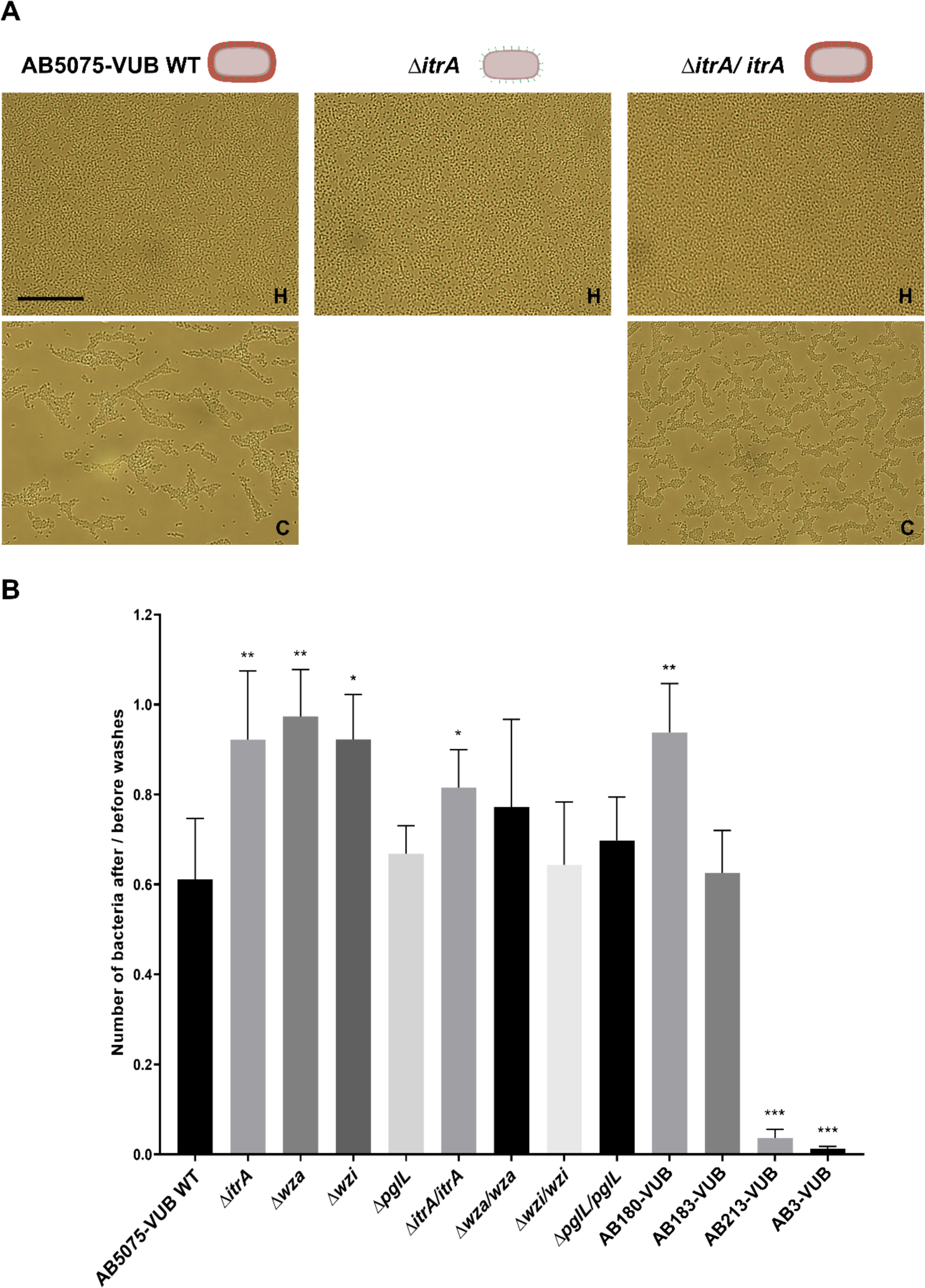
*A. baumannii* adhesion to polystyrene. (A) Representative microscopy pictures of the adhesion assay for AB5075-VUB WT, *ΔitrA,* and *ΔitrA/itrA.* Capsulated strains present two different patterns after washes. They are indicated in the pictures: homogeneous distribution of bacteria on the surface: H, and bacteria clustered on the surface: C. Scale bar: 50 µm (B) The ratio of the estimated number of cells after wash/ before washing for each strain. A one-way ANOVA was performed for each set of experiments and showed significant differences among means. Then, unpaired student tests (t-tests) were performed on the data by comparing each strain to the AB5075-VUB WT (reference). The difference is significant if the P-value (P) <0.05. GP(GraphPad): 0.0332<P-value<0.05 (*), 0.0021<p-value<0.0332 (**), 0.0002<p-value<0.0021 (***), p-value<0.0001(****). Statistics were performed on the average fold change value for each tested strain and compared to the average fold change obtained with the AB5075-VUB WT strain.

### The presence of the *A. baumannii* capsule impacts natural transformation frequency

While performing routine transformation using the natural competence of *A. baumannii* with the AB5075 strain, we observed a systematically lower number of transformants with non-capsulated parental strains than when we used capsulated ones. To confirm this observation, we calculated transformation frequencies using a set of capsule mutants. We investigated the capsule’s impact in the natural transformation process with both linear DNA and a plasmid using AB5075 WT, *itrA::ISAba13*, the CO revertant originating from this strain, and the capsule mutants generated so far. Using overlap extension PCR, we generated a linear DNA fragment containing an upstream and downstream homologous region with the Tn7 site and an *aaC* (encoding for apramycin resistance) cassette in between to select transformants: *the attTn7::aaC* fragment. We also used the pASG1-sfGFP plasmid carrying *sfGFP* and an *aaC* cassette for the selection of the transformants. The bacteria and DNA were incubated for 6 hours on the motility medium and then plated on LB-agar and LB-agar with 30µg/mL apramycin to obtain the total number of bacteria and the number of transformants. The calculated frequencies obtained in these experiments are presented in **Figure 5**.

**Figure 5:**
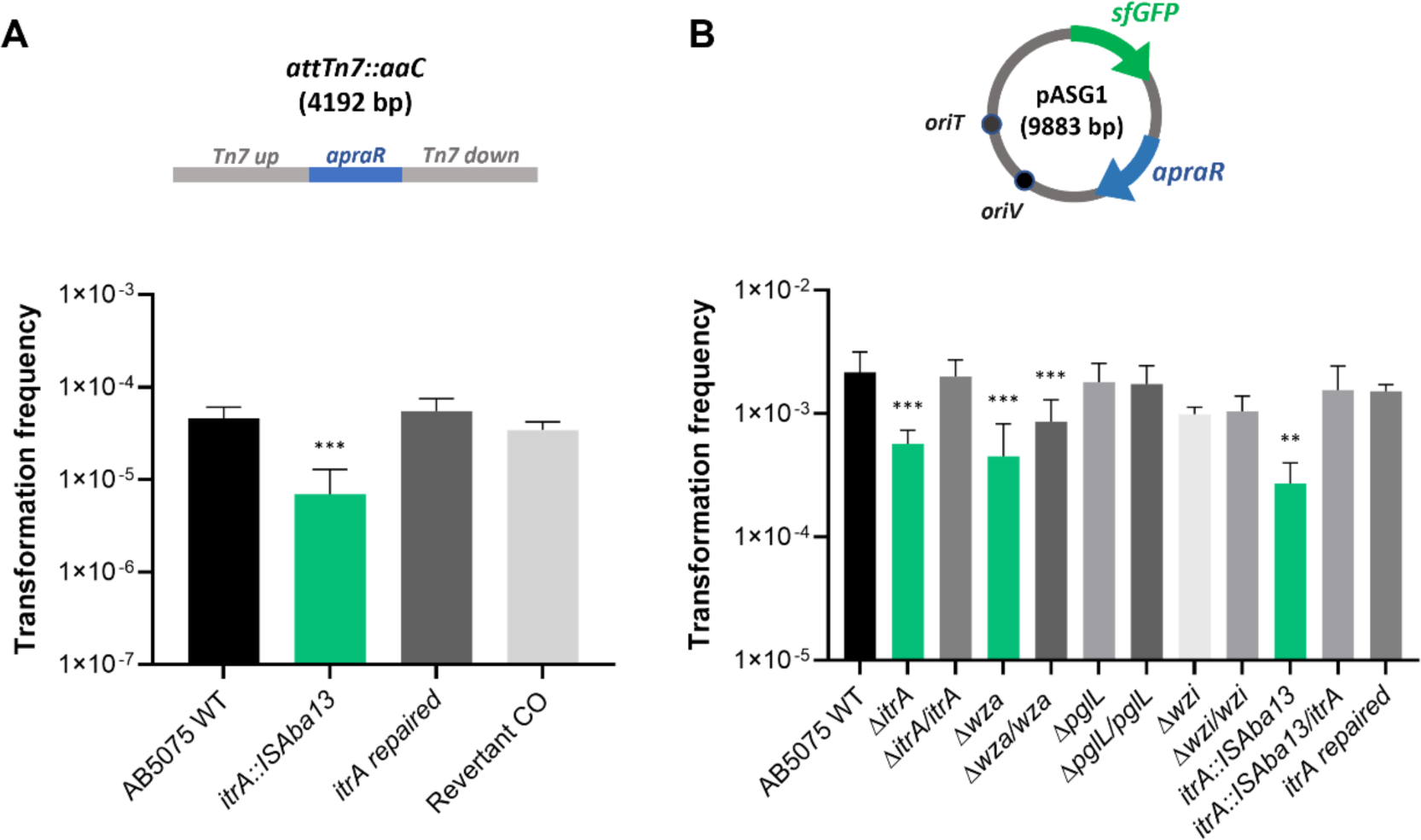
Natural transformation frequencies of the capsule mutants. (A) Frequencies were obtained using linear DNA (att*Tn7::aaC*) and (B) the pASG1-sfGFP plasmid. Three independent biological replicates (n=3) were performed with all the strains. Non-capsulated strains are represented in green. A one-way ANOVA was performed for each set of experiments and showed significant differences among the means. Then unpaired student tests (t-tests) were performed on the data by comparing each strain to the AB5075-VUB WT (reference). The difference is significant if the P-value (P) <0.05. GP(GraphPad): 0.0332<P-value<0.05 (*), 0.0021<p-value<0.0332 (**), 0.0002<p-value<0.0021 (***), p-value<0.0001(****).

A 3 to 5-fold decrease in the transformation frequency was observed for the non-capsulated mutant *itrA::ISAba13* with linear DNA compared to the WT. A similar difference was observed for the non-capsulated mutants *itrA::ISAba13*, Δ*itrA,* and *Δwza* with the pASG1-sfGFP plasmid. No significant difference in the average frequency was observed for the *Δwzi* and *ΔpglL* mutants compared to the WT strain in the tested conditions. The frequencies of the complementation strains are restored to the WT level except for the *Δwza/wza* complementation, for which the frequency was only partially restored.

In this experiment, we observed a decreased natural transformation frequency for the non-capsulated strains of AB5075 compared to the capsulated ones. Before that, a positive or negative correlation between capsule production and competence was, to the best of our knowledge, never reported in *A. baumannii*.

## Discussion

Adhesion is a complex and multifactorial process, which represents a crucial step in host cell colonization and the formation of stable biofilms (16,17). However, polysaccharide capsules can form a barrier to such interactions. Several studies with diverse bacterial models have reported that capsule production represents a trade-off and can impair adhesion, auto-aggregation, host-cell invasion, and biofilm formation by masking appendages such as fimbriae and adhesins, or through its viscosity/repulsive ability (18–22). However, to our knowledge, our report represents one of the first studies directly linking capsule production and adhesion in *A. baumannii*. A recent study of experimental evolution with *A. baumannii* supports our results: they observed that thinner capsules increased adhesion and internalization by macrophages and that enhanced capsular exopolysaccharide chain length led to the opposite phenotypes (23).

In our study, we generated and characterized capsule mutants of AB5075-VUB. Deletion of *wza* leads to loss of capsule production and virulence. We also identified a growth defect in the stationary phase. A recent report showed that blockages in late steps of the Wzy-dependent pathway are lethal in the ATCC17978 *A. baumannii* strain unless suppressed by mutations in the ItrA or BfmS-dependent pathways, but could be strain-dependent (15). Such modifications have also been shown to lead to toxicity and cell wall defects in *K. pneumoniae,* through possible accumulation of Und-P or capsule intermediates (24,25). Care should be taken about capsule mutants used in phenotypic studies, especially if looking at virulence. Consequently, we used the *ΔitrA and ΔpglL* strains in parallel with the *Δwza* strain to ensure the adhesion phenotype observed is linked to the absence of the capsule. Deletion of *ΔpglL* does not impact capsule production and virulence in the tested condition. Deletion of *wzi* leads to an intermediate capsule production that is probably transient. We showed that the *Δwzi* is not virulent in *G. mellonella* but impacts the fitness of the larvae (partial melanization). Sanchez-Larrayoz *et al*., 2017 reported a decreased virulence of a *wzi* mutant in human serum, supporting our observation (26). Moreover, in a study by Tickner *et al*., 2021, Wzi was shown to be required for the retention of CPS on the cell surface of AB5075, and an increased proportion was found outside of the cell for the *wzi::T26* mutant (10). An increased ratio of CPS secreted in the hemolymph by our *Δwzi* mutant could potentially lead to an immune response of the larvae without killing them and explains the partial melanization of the *G. mellonella* larvae.

We included some isolates from a previous study: one low-capsulated (AB180-VUB), one with an intermediate/low capsulation (AB183-VUB), and two constitutively mucoid (AB3-VUB and AB213-VUB) displaying thick capsules (2). AB180-VUB was shown to be avirulent in *G. mellonella*, AB183-VUB weakly virulent, and surprisingly, AB3-VUB was weakly virulent when AB213-VUB was fully virulent, even though both strains carry the same capsule type: KL40. However, TEM pictures showed that the AB3-VUB capsule is much looser than AB213-VUB (2), which stands firmly around the bacterial cells. The capsule size and related structure are very heterogeneous among clinical isolates (2). This observation partially originates from the several K-locus types, but other factors outside of the K-locus can influence these phenotypes (27). KL-type alone does not always represent the capsule phenotype and highlights the importance of phenotypical characterization.

Interestingly, pellet size after centrifugation correlates with the capsulation level, and their stability gave us a first clue about the ability of *A. baumannii* cells to adhere to each other. We saw that non/low-capsulated strains form much smaller and more stable pellets than capsulated ones. The constitutively mucoid AB3-VUB pellet is highly unstable and correlates with a low adhesion level. In other studies, auto-aggregation was shown to be decreased between capsulated bacteria compared to non-capsulated ones (28,29). The assessment of the aggregation of bacterial cells to each other depending on the capsulation of *A. baumannii* will be relevant since it is a critical step in biofilm formation.

We then showed that the capsule of AB5075-VUB impairs adhesion to epithelial cells and impacts adhesion to a polystyrene surface. However, in a previous study, deletion of *wza* in the SKLX024256 clinical isolate was shown to impair adhesion to A549 epithelial cells (9), whereas we see the opposite with the *Δwza* with AB5075 and in our experimental conditions. This could be linked to using different experimental settings, strain-dependent factors influencing adhesion or potential consequences of deletion of a late-stage component of the Wzy-dependent pathway that seems to be more or less tolerated by *A. baumannii* strains (8,15). However, we used deletions of *itrA* and *pglL* to show that the observed adhesion phenotype is related to capsule production. In agreement with our observations, the influence of *A. baumannii*’s capsule on adhesion and natural transformation frequency was also recently reported (BIORXIV-2024-580519v1-Cooper *et al.,* 2024).

Moreover, the low-capsulated AB180-VUB isolate is also more adherent to epithelial cells and consistently adherent to polystyrene. The AB183-VUB and *Δwzi* strains depicted intermediate adhesion ability to epithelial cells, but *Δwzi* was more adherent to the polystyrene in the tested conditions. Finally, we showed that mucoid capsules, especially the loose capsule of AB3-VUB, seem to impair adhesion to epithelial cells and polystyrene. This does not exclude that different conditions and surfaces could change the observed phenotypes and need to be investigated.

Recent reports describe and support the facultative intracellular lifestyle of *A. baumannii* (30). Indeed, some isolates have been shown to survive and even proliferate inside eukaryotic cells (31) and macrophages (32). According to our results, most isolates of *A. baumannii* probably require a lower level of capsulation to enter the host cells. The capsule size, structure, and composition likely modulate this intracellular lifestyle. On the other hand, even if strong capsulation inhibits adhesion, it remains a significant virulence and resistance factor for *A. baumannii* to survive the host immune defenses, antimicrobial, disinfectants, and desiccation (8,13). Some studies report that thickened capsules lead to reduced biofilm formation, some that capsule is involved in the process, and some did not identify any correlation (9,11,14,23). The relationship between capsule production and biofilm formation is complex, and it is difficult to determine which influences the other. The amount of capsules produced may vary depending on the growth phase, environmental conditions, and potential stresses (33,34), but little is known about its regulation in *A. baumannii*. Production of the capsule represents a trade-off, and identifying such phenotypes could explain the role of both VIR-O and AV-T in variant in phase variation (35) in modulating the resistance and virulence of *A. baumannii*. However, more research needs to be done on its regulation and how it is synchronized with the expression of other surface-exposed proteins involved in adhesion.

Here, we showed that deletions of genes involved in the biosynthesis and export of the capsule impact the natural transformation process in AB5075 in the tested conditions. Further investigations would be needed to determine which step(s) of the natural transformation is impacted by the presence/absence of the *A. baumannii* capsule.

## Conclusion

In our study, we assessed the effects of capsule production on adhesion to epithelial cells and abiotic polystyrene surface of different mutants from the AB5075-VUB strains and 4 clinical isolates with distinct capsulation levels. We showed that thick capsule and mucoid phenotype impair adhesion, whereas capsule loss increases it drastically in the tested conditions. This study highlights the critical role of *A. baumannii* capsule regulation in adhesion and the diversity of phenotypes found among isolates. The bacterial capsule can protect or sensitize against chemical compounds, environmental hazards, and bacteriophages. This illustrates the importance of the trade-off of capsule production in *A. baumannii* for its virulence, natural competence, environmental, and hospital persistence. A better understanding of adhesion to biotic and abiotic surfaces will help understand *A. baumannii’s* success in clinical settings and design new antimicrobial strategies such as novel anti-virulence and anti-resistance compounds.

## First’s author biography

Dr. Clémence Whiteway graduated from the Pasteur Institute. She did her PhD in the Van der Henst’s Lab at the Vrije Universiteit Brussels / Flanders Institute for Biotechnology.

## Supplementary materials

**Table S1.** Strains and plasmids.

| Name of the strain | Genotype | Reference |
| --- | --- | --- |
| AB5075-VUB WT | AB5075-VUB (CP070362)<br>BioProject - PRJNA701627 – KL25 | (Whiteway *et al.*, 2021) |
| $\Delta itrA::sacB-aaC$ | $\Delta itrA::sacB-aaC$ intermediate strain to generate *itrA* deletion in the AB5075-VUB WT genome | (Whiteway *et al.*, 2021) |
| $\Delta itrA$ | $\Delta itrA$ | (Whiteway *et al.*, 2021) |
| $\Delta itrA/sacB-aaC$ | $\Delta itrA$ Tn7::*Pst-sacB-aaC* intermediate strain to clone *itrA* at the Tn7 site | (Whiteway *et al.*, 2021) |
| $\Delta itrA/itrA$ | $\Delta itrA$ Tn7::*Pst-itrA* | (Whiteway *et al.*, 2021) |
| $\Delta wza::sacB-aaC$ | $\Delta wza::sacB-aaC$ intermediate strain to generate *wza* deletion in the AB5075-VUB WT genome | This study |
| $\Delta wza$ | $\Delta wza$ | This study |
| $\Delta wza/sacB-aaC$ | $\Delta wza$ Tn7::*Pst-sacB-aaC* intermediate strain to clone *wza* at the Tn7 site of $\Delta wza$ | This study |
| $\Delta wza/wza$ | $\Delta wza$ Tn7::*Pst-wza* | This study |
| $\Delta wzi::sacB-aaaC$ | $\Delta wzi::sacB-aaaC$ intermediate strain to generate *wzi* deletion in the AB5075-VUB WT genome | This study |
| $\Delta wzi$ | $\Delta wzi$ | This study |
| $\Delta wzi/sacB-aaC$ | $\Delta wzi$ Tn7::*Pst-sacB-aaC* intermediate strain to clone *wzi* at the Tn7 site of $\Delta wzi$ | This study |
| $\Delta wzi/wzi$ | $\Delta wzi$ Tn7::*Pst-wzi* | This study |
| $\Delta pglL::sacB-aaC$ | $\Delta pglL::sacB-aaC$ intermediate strain to generate *pglL* deletion in the AB5075-VUB WT genome | This study |
| *ΔpglL* | *ΔpglL* | This study |
| *ΔpglL/sacB-aaC* | *ΔpglL Tn7::Pst-sacB-aaC* intermediate strain to clone *pglL* at the Tn7 site of *ΔpglL* | This study |
| *ΔpglL/pglL* | *ΔpglL Tn7::Pst-pglL* | This study |
| AB3-VUB | AB3-VUB (CP091378) KL40 | (Valcek et al., 2023; Valcek, Nesporova, et al., 2022) |
| AB180-VUB | AB180-VUB (CP091359) KL3 | (Valcek et al., 2023; Valcek, Nesporova, et al., 2022) |
| AB183-VUB | AB183-VUB (CP091357) KL9 | (Valcek et al., 2023; Valcek, Nesporova, et al., 2022) |
| AB213-VUB | AB213-VUB (CP091349) KL40 | (Valcek et al., 2023; Valcek, Nesporova, et al., 2022) |
| **Plasmids** |  |  |
| pASG1-1 | Pst-sfGFP for constitutive and strong expression of cloned genes and *sfGFP* cloning | (Godeux *et al.*, 2018) |
| pMHL2 | Apramycin resistance (*aac*) and sucrose sensitivity (*sacB*) | (Godeux *et al.*, 2018) |

**Tables S2.**
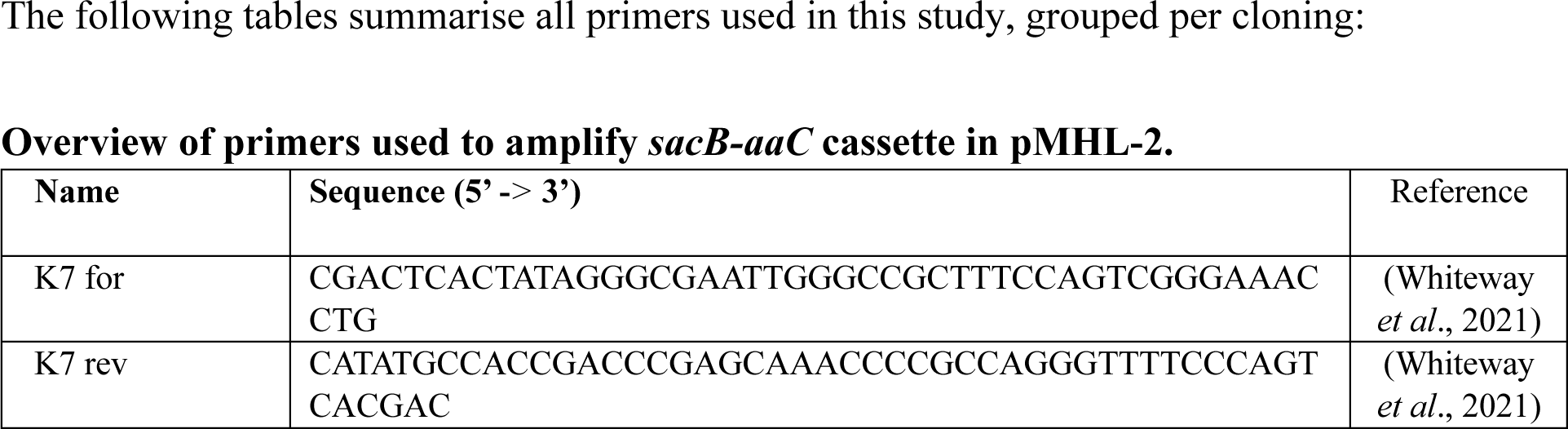
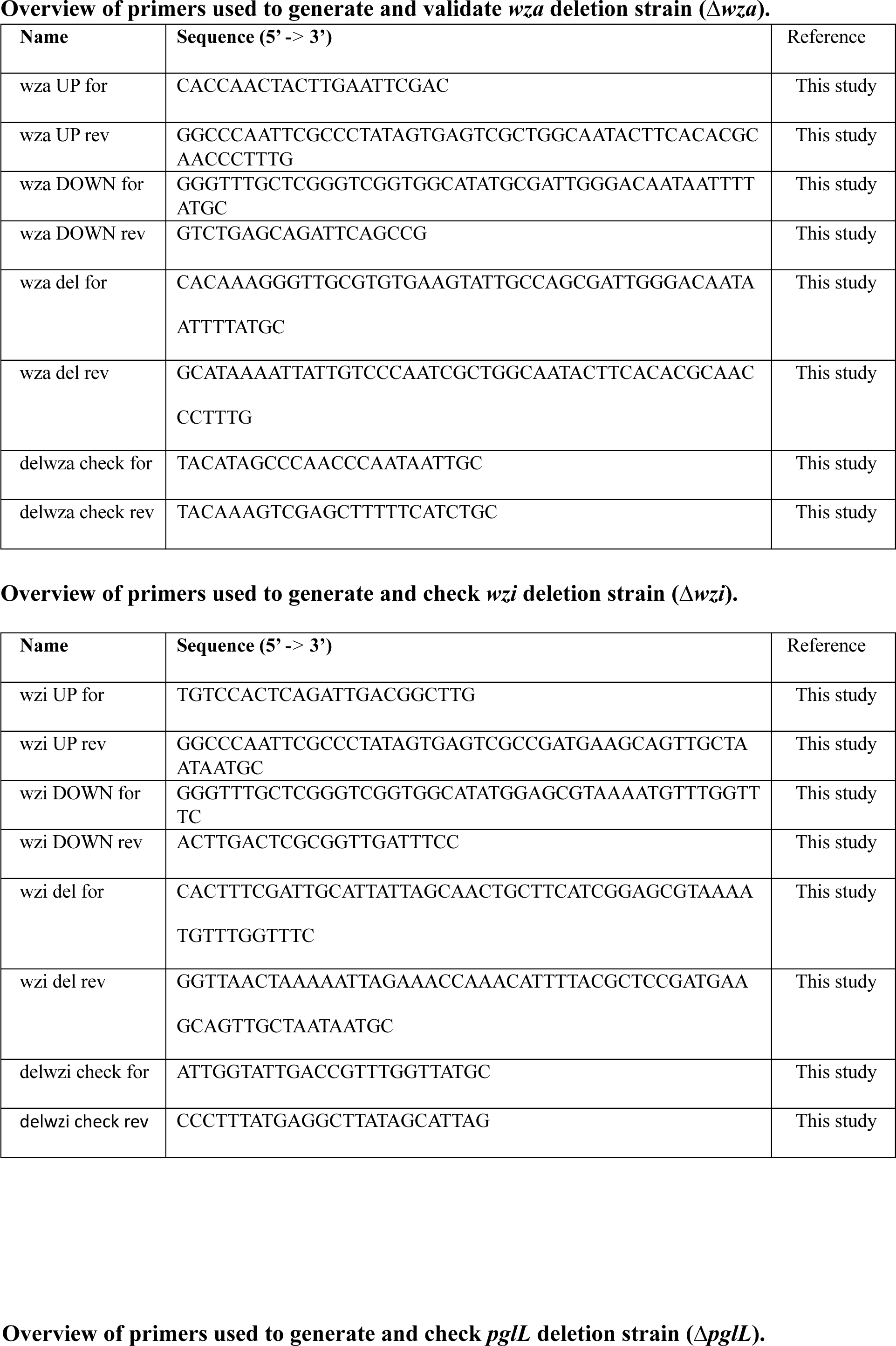
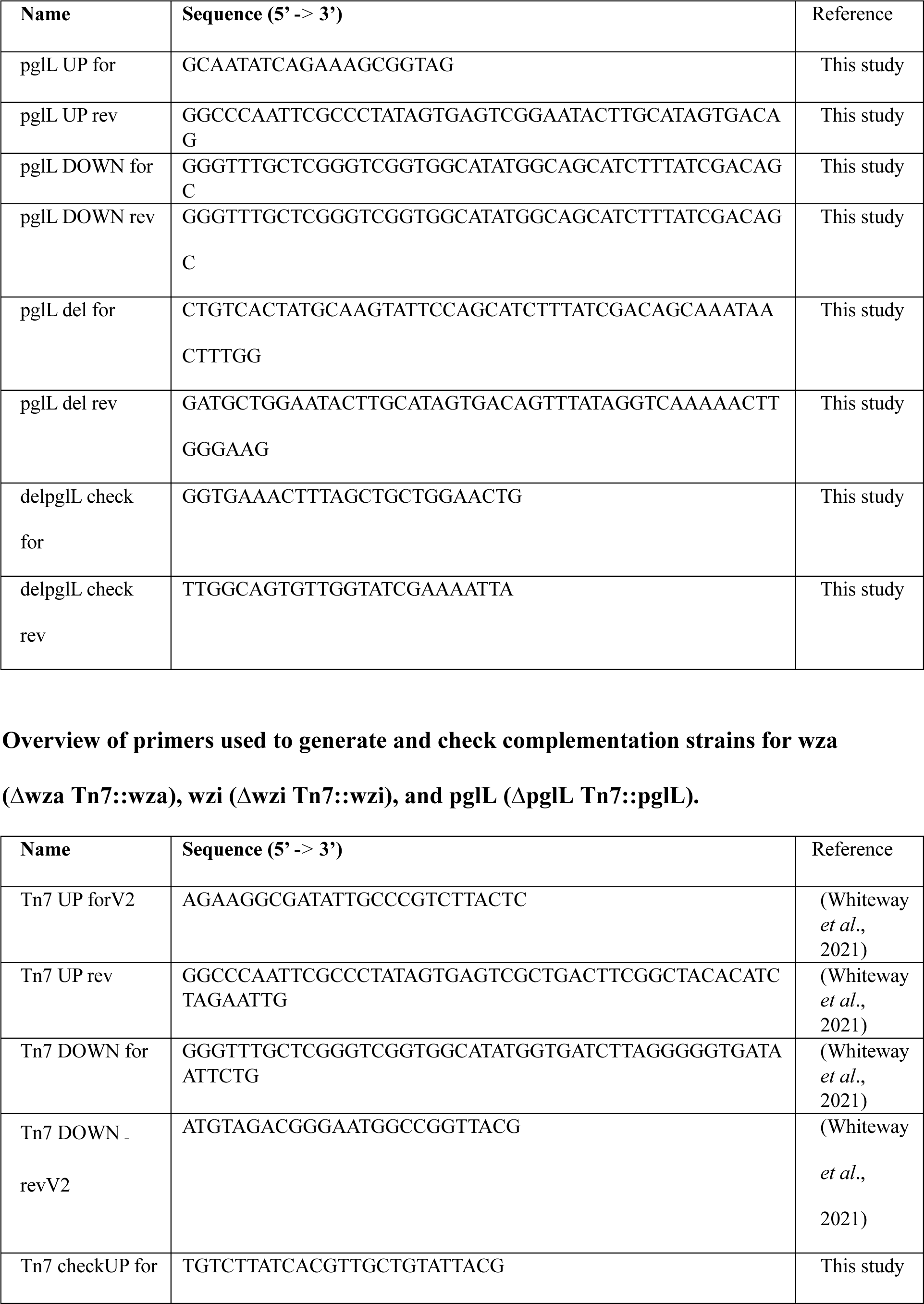
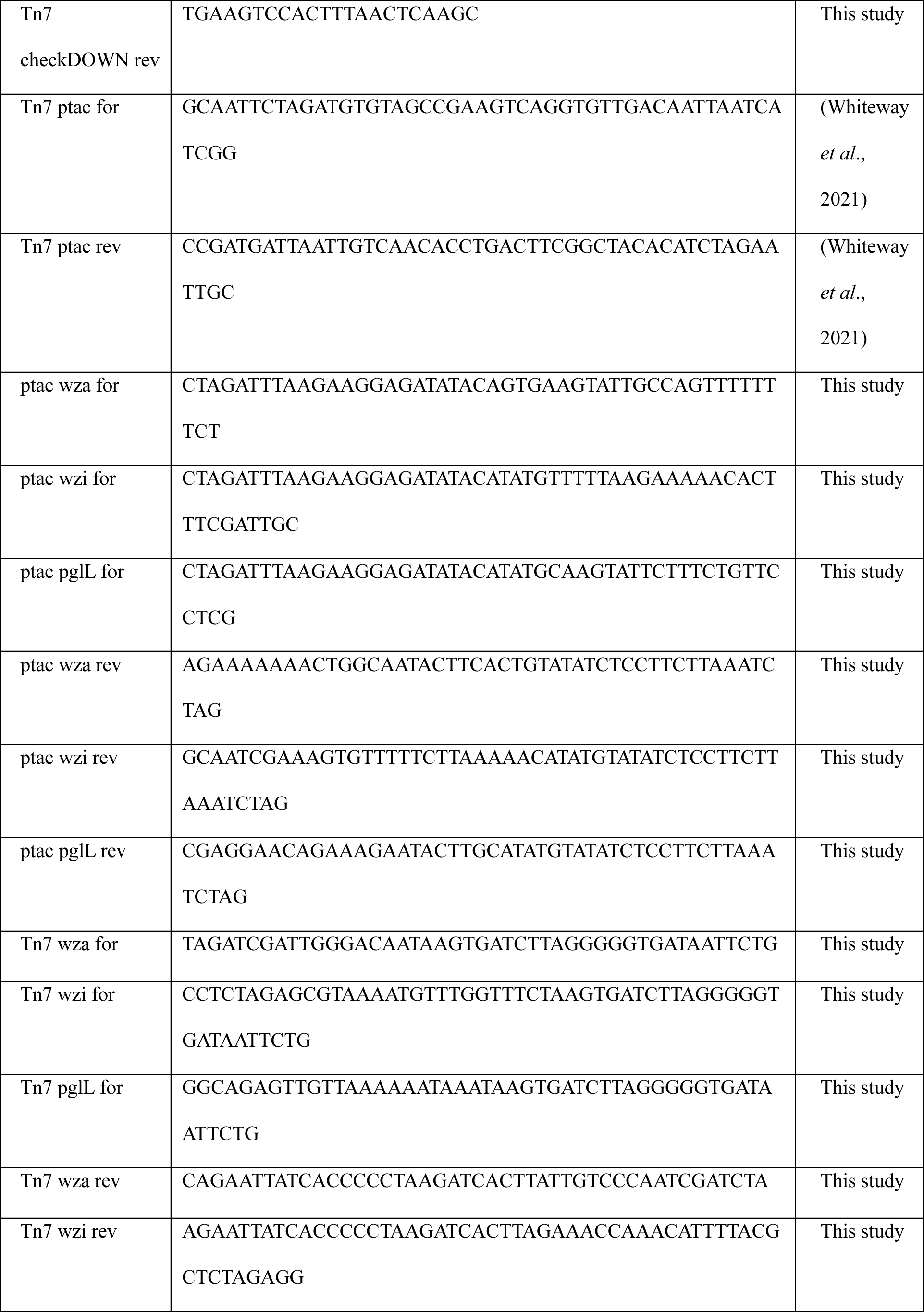
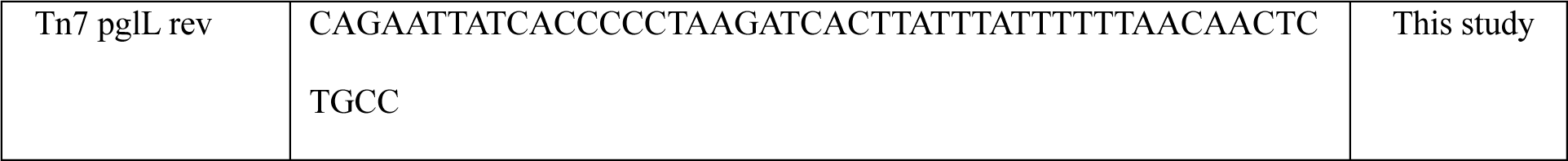
List of primers used in this study.

**Figure S1.**
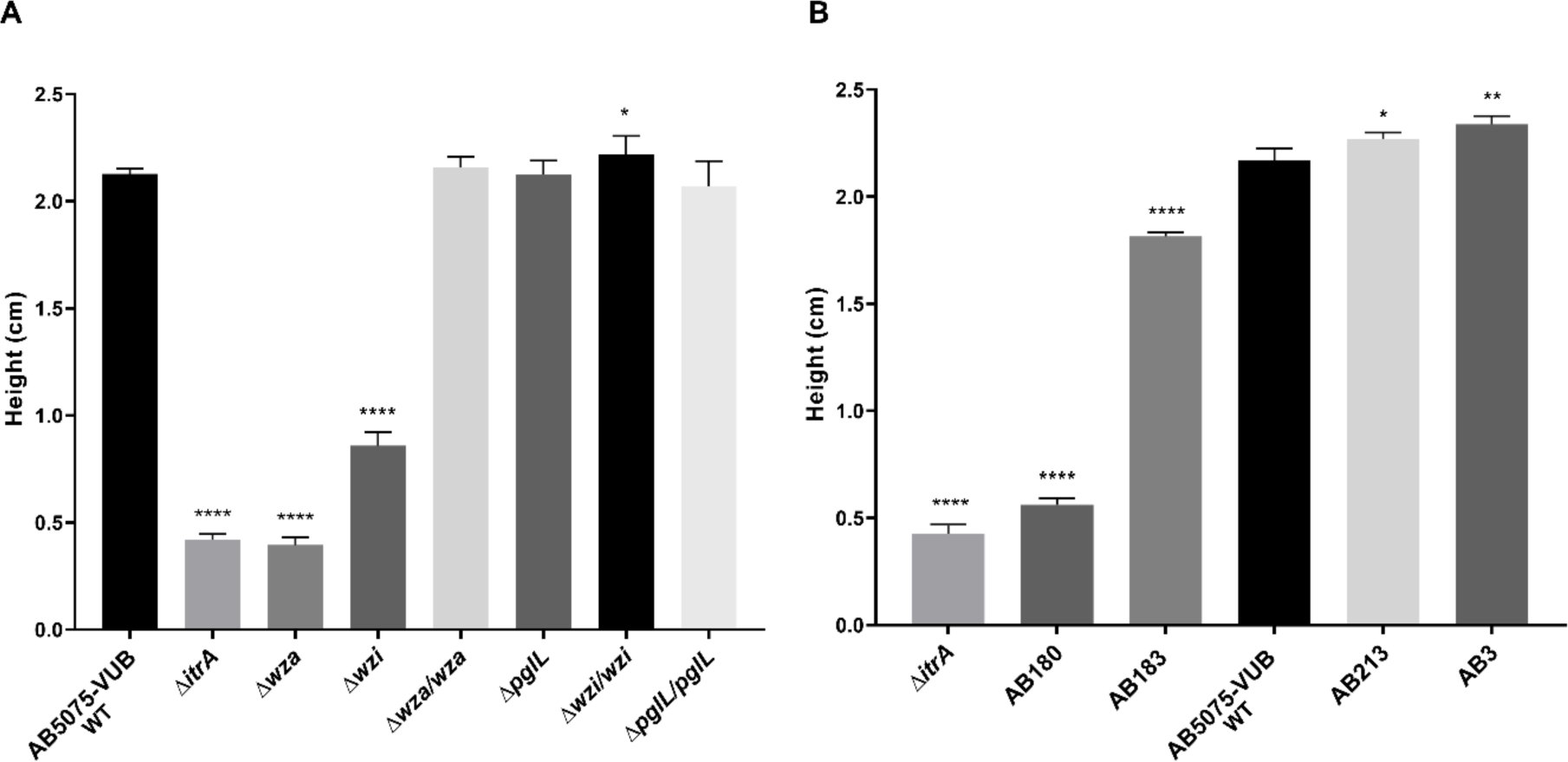
Semi-quantification of capsule production using density gradients. A) Average height of bands after centrifugation in the density gradients for capsule and *O*-linked glycosylation mutants of AB5075-VUB. B) Average height of bands after centrifugation in the density gradients for the clinical isolates. The height of the band (cm) was measured from the bottom of the tube (y-axis) for the different strains (x-axis). The results were compared using a one-way ANOVA and showed a significant difference among means. Then unpaired student tests (t-tests) were performed on the data by comparing each strain to the WT (reference The difference is significant if the P-value (P) <0.05. GP(GraphPad): 0.0332<P-value<0.05 (*), 0.0021<p-value<0.0332 (**), 0.0002<p-value<0.0021 (***), p-value<0.0001(****). The *ΔitrA* mutant was used as a control for the absence of capsule production. This experiment was performed in a biological quadruplicate.

**Figure S2.**
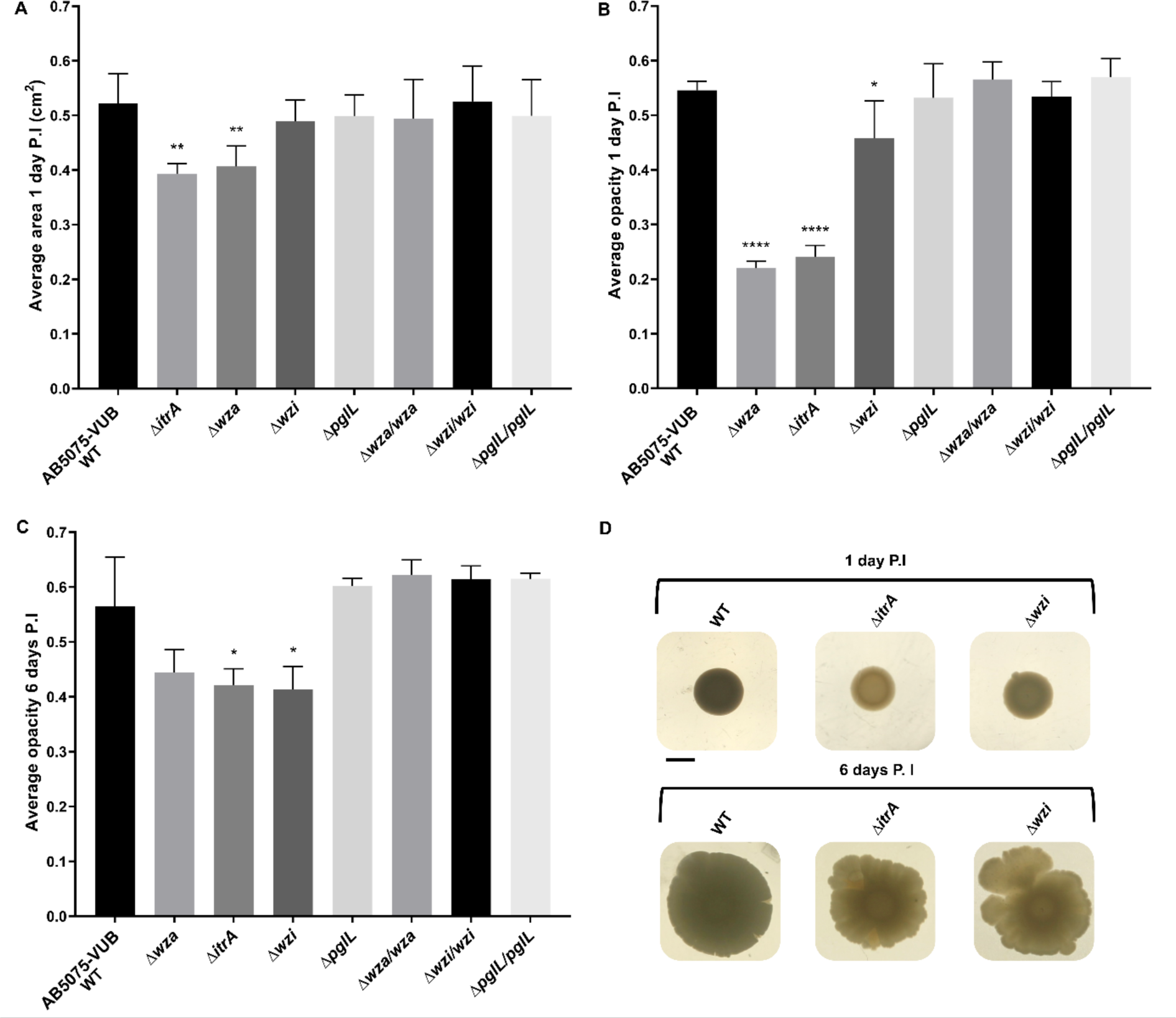
Area and opacity of macrocolonies on solid media after 24h at 37°C. A) Average areas of the macrocolonies after 24h post-inoculation (P.I) on solid LB-agar media were measured using ImageJ. B) Average opacities after 1 day P.I and C) 6 days P.I was measured using ImageJ (see Material and Methods). One-way ANOVA was performed and showed significant differences among means. Then, unpaired student tests (t-tests) were performed on the data by comparing each strain to the WT (reference). The difference is significant if the P-value (P) <0.05. GP(GraphPad): 0.0332<P-value<0.05 (*), 0.0021<p-value<0.0332 (**), 0.0002<p-value<0.0021 (***), p-value<0.0001(****). D) Pictures of the macrocolonies 1 day P.I and 6 days P.I. Scale bar: 0.5 cm.

**Figure S3.**
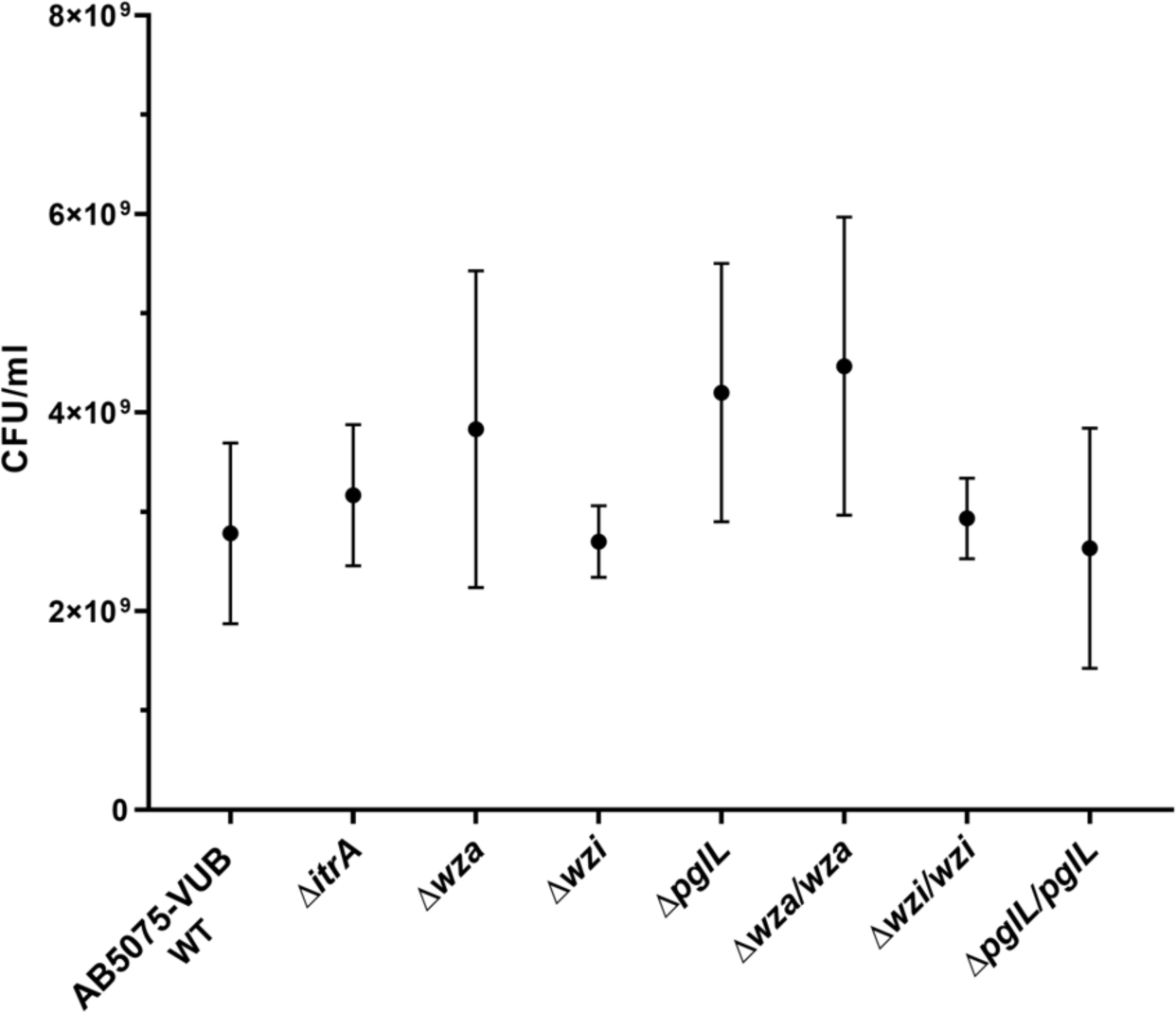
Colony forming units (CFU/ml) of AB5075-VUB and mutants. Bacterial concentrations were obtained by plating a stationary phase overnight culture of the AB5075 WT and mutants (CFU/ml). Bacterial cultures were grown for 16h at 37°c). The results were compared using a one-way ANOVA and showed no difference among means: (P-value= 0,3061 and F= 1.313).

**Figure S4.**
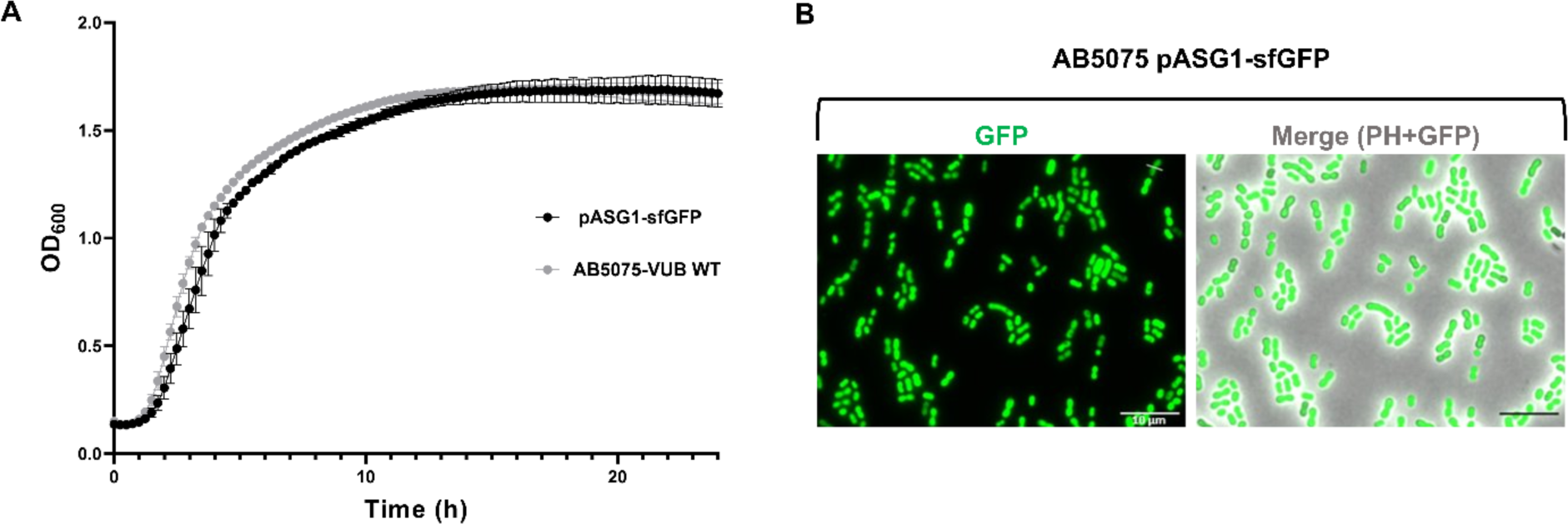
Validation of the fluorescence of the AB5075 pASG1-sfGFP strain. A) Growth measurements of AB5075 WT and the same strain carrying the pASG1-sfGFP plasmid (OD_600_ measurements over 24h). They have similar growth in the tested conditions. B) Microscopy pictures of the AB5075 strain carrying the pASG1-sfGFP plasmid. The percentage of GFP+ bacteria was determined, 99,6% of 3837 bacteria were GFP+ (SD=4.429. 10^−5^). Scale bar: 10 µM.

**Figure S5.**
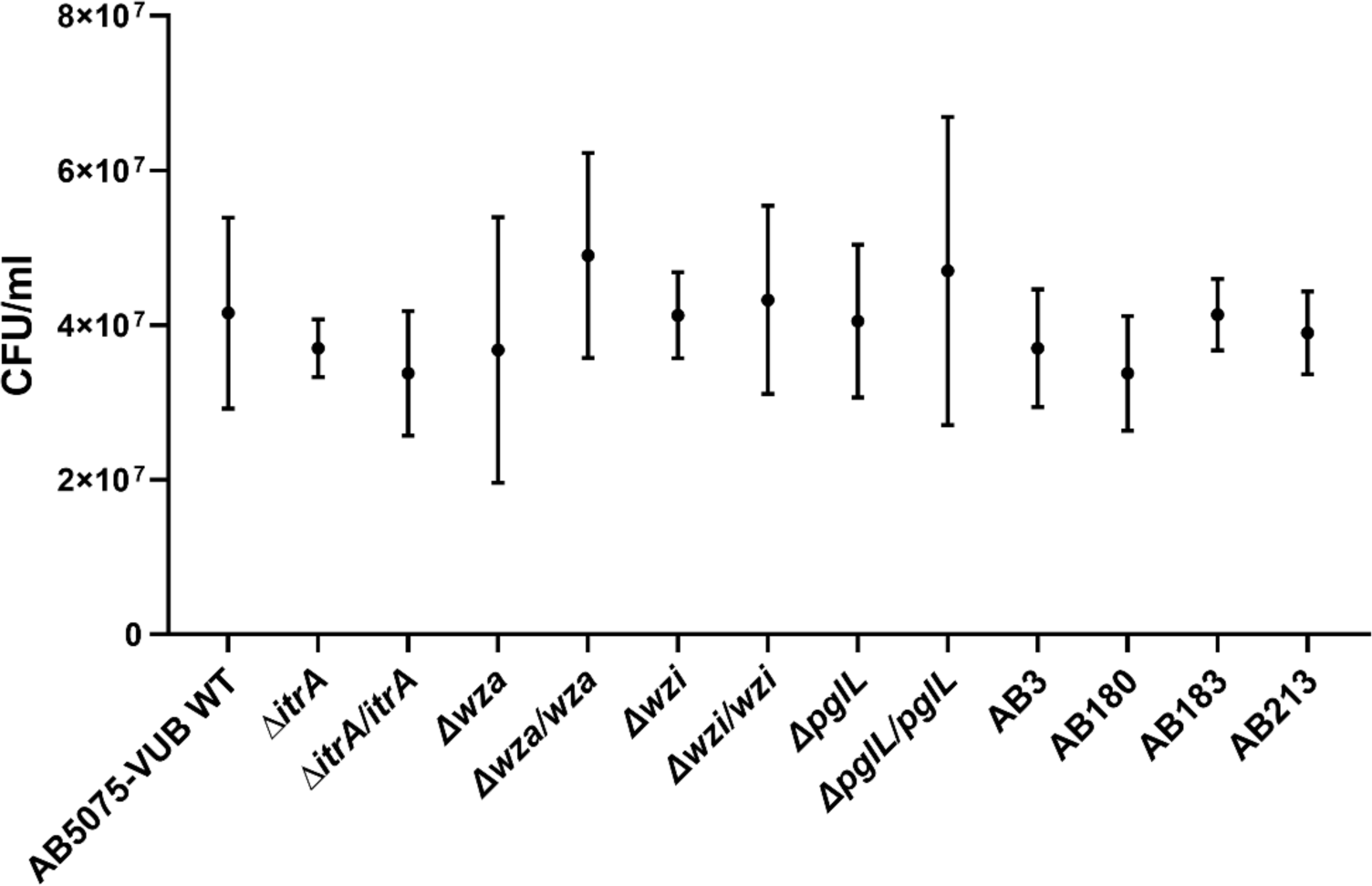
Colony forming units (CFU/ml) of all the inoculums before A549 epithelial cells adhesion assay. The results were compared using a one-way ANOVA and showed no difference among means: (P-value= 0,3061 and F= 0,6857).

**Figure S6.**
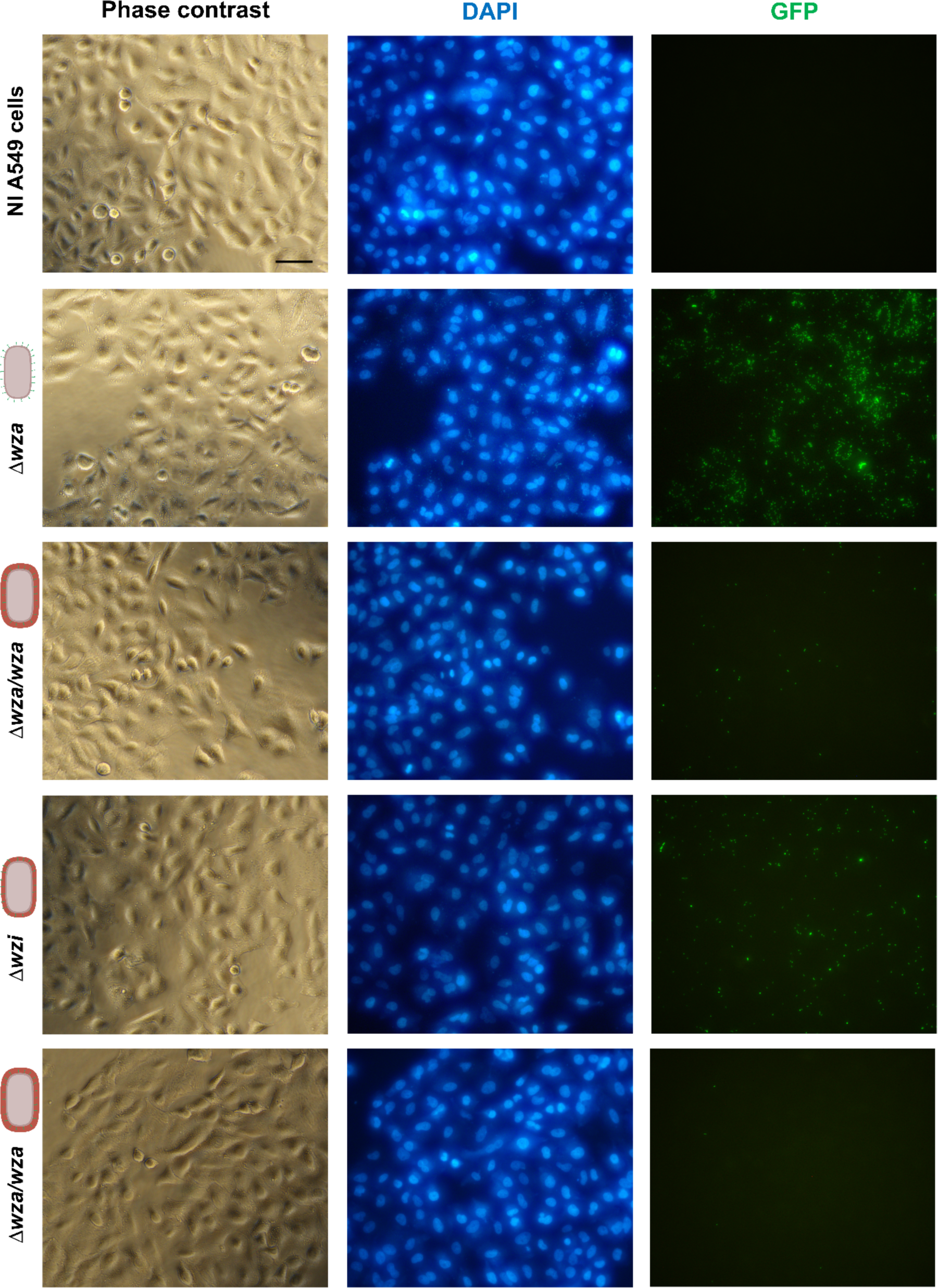
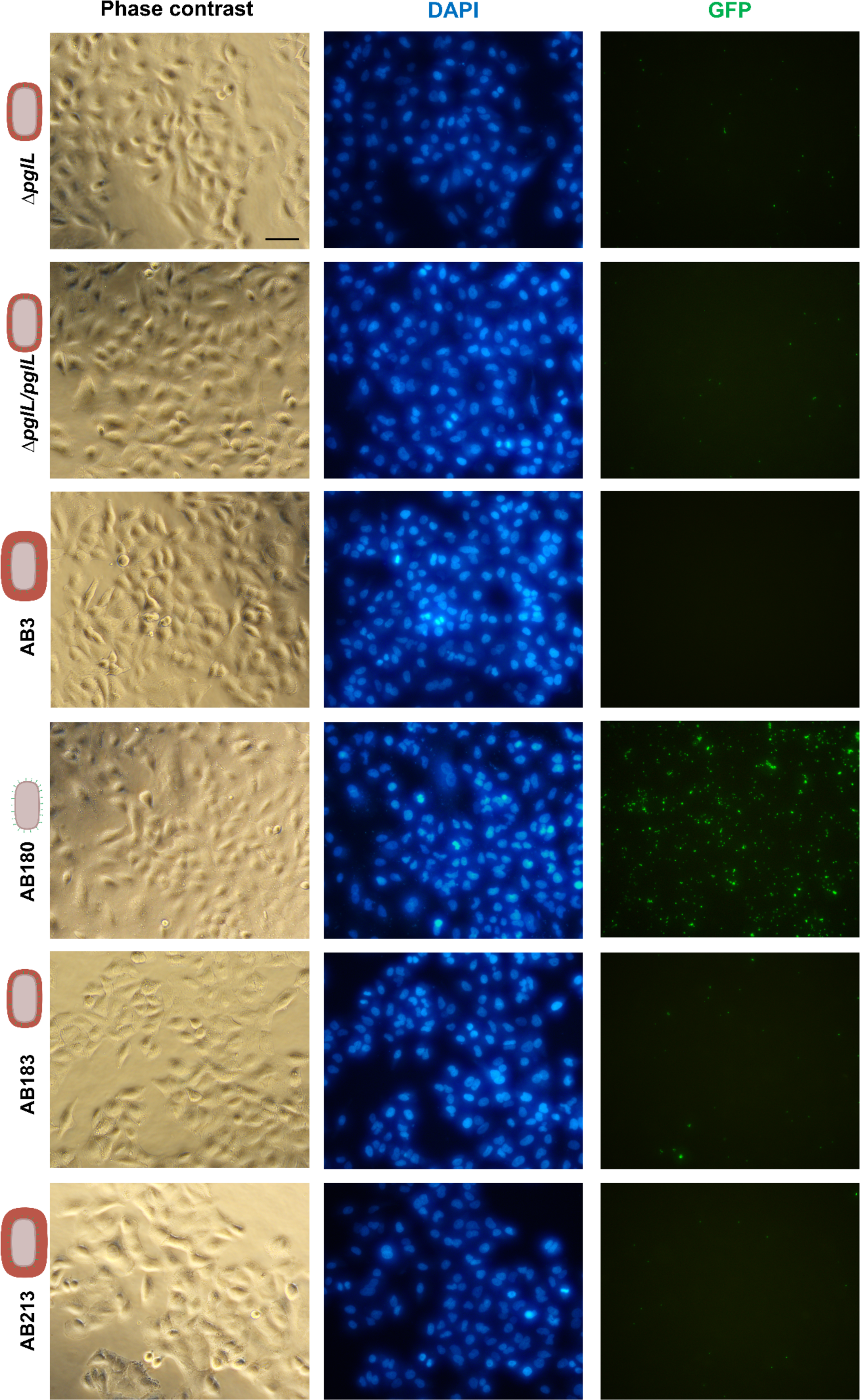
Adhesion of *A. baumannii* to epithelial cells. Microscopy pictures representing the different adhesion profiles observed after adhesion assay on A549 epithelial cells. Scale bar: 50 µm.

**Figure S7.**
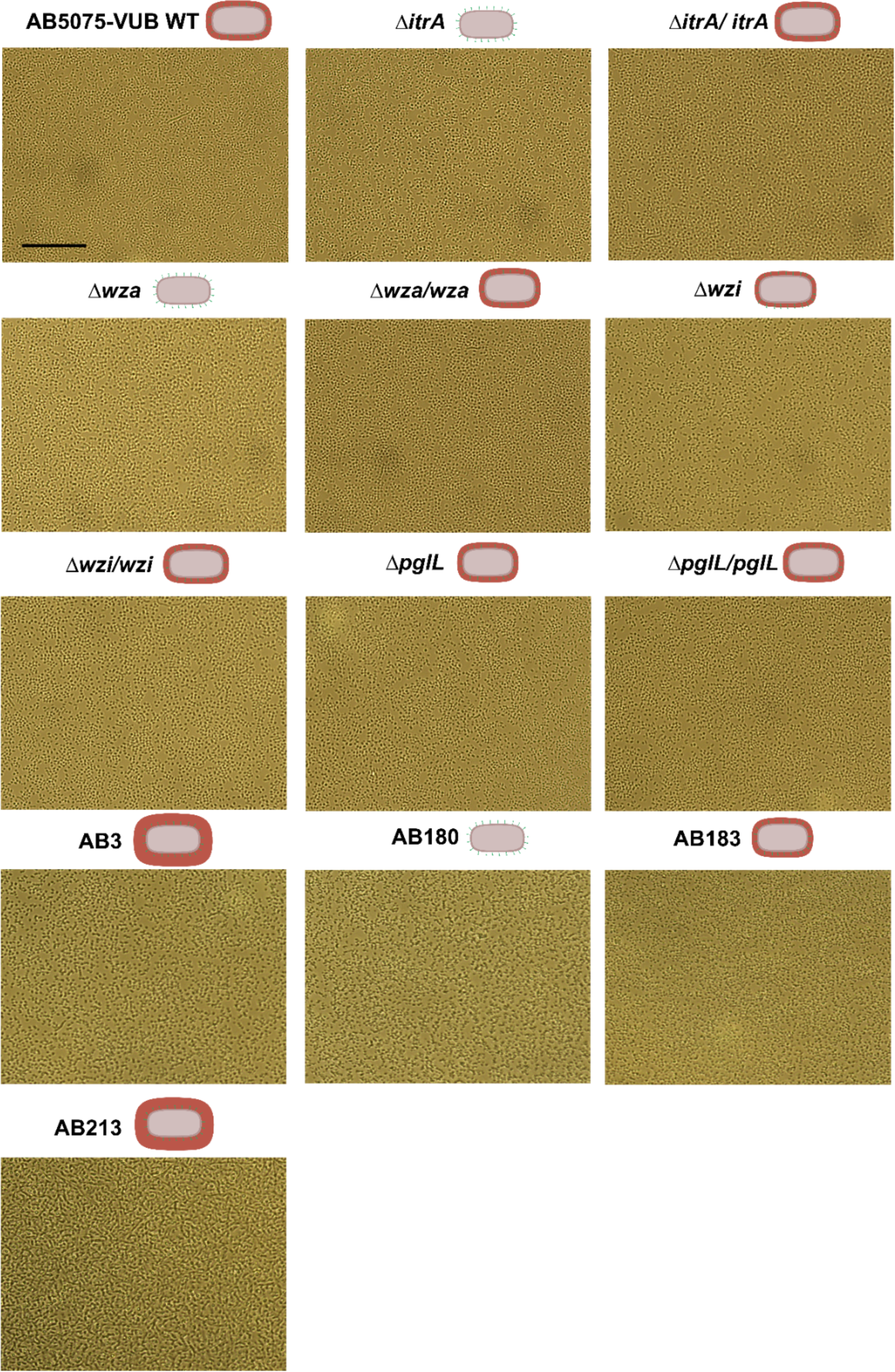
Adhesion of *A. baumannii* to polystyrene (before washing). Microscopy pictures representing the different adhesion profiles observed before adhesion on a non-treated 96-well plate. Scale bar: 50 µm.

**Figure S8.**
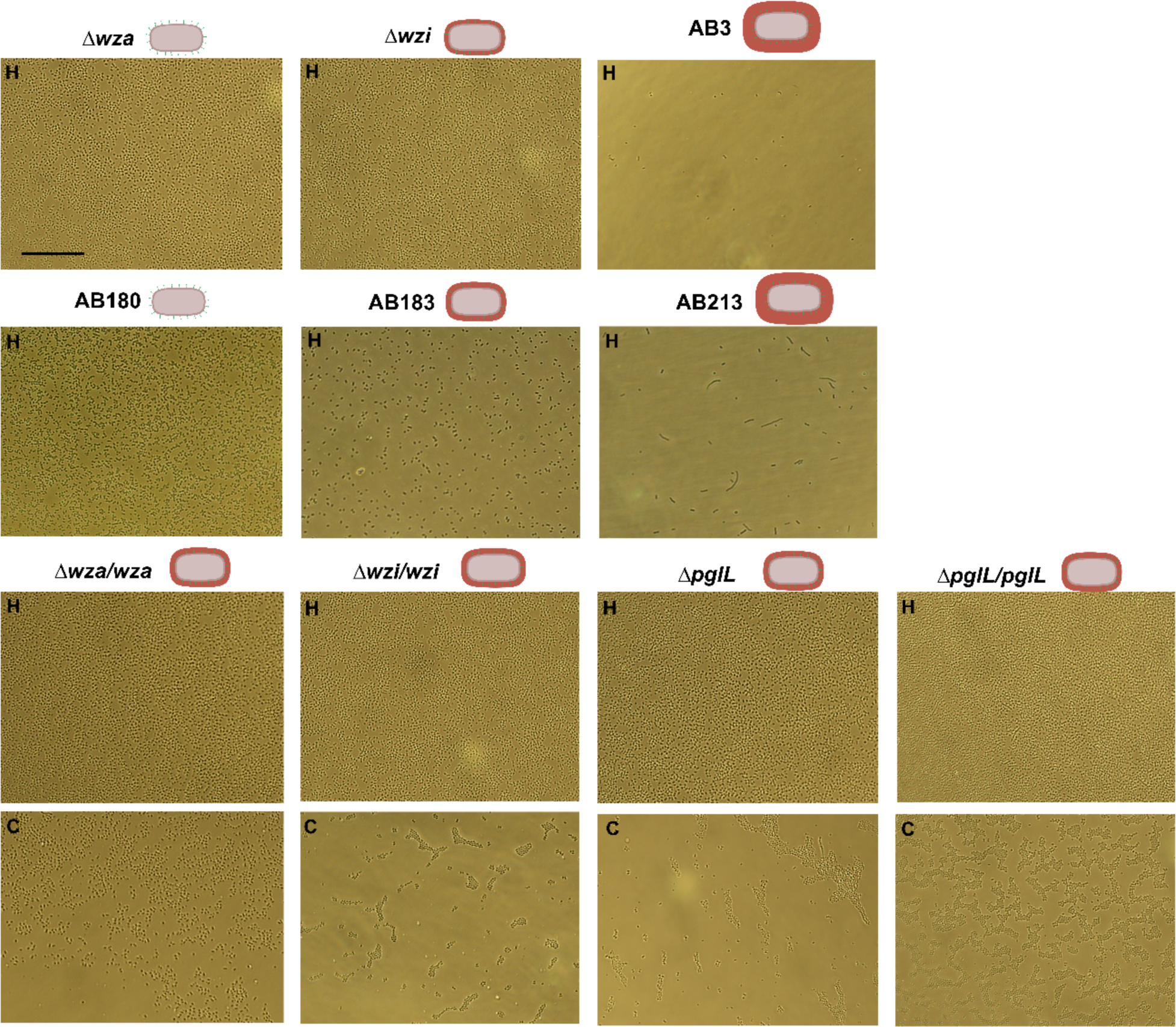
Adhesion of *A. baumannii* to polystyrene (after washing). Microscopy pictures representing the different adhesion profiles observed before adhesion on a non-treated 96-well plate. Scale bar: 50 µm.

## References

1. Jacobs AC, Thompson MG, Black CC, Kessler JL, Clark LP, McQueary CN, et al. AB5075, a highly virulent isolate of *Acinetobacter baumannii*, as a model strain for the evaluation of pathogenesis and antimicrobial treatments. MBio. 2014;5(3):1–10.

2. Valcek A, Philippe C, Whiteway C, Robino E, Nesporova K, Bové M, et al. Phenotypic Characterization and Heterogeneity among Modern Clinical Isolates of *Acinetobacter baumannii*. Microbiol Spectr. 2023;11(1).

3. Whiteway C, Valcek A, Philippe C, Strazisar M, De Pooter T, Mateus I, et al. Scarless excision of an insertion sequence restores capsule production and virulence in *Acinetobacter baumannii*. ISME J. 2021;(August 2021):1–5.

4. Kumar A, Dalton C, Cortez-Cordova J, Schweizer HP. Mini-Tn7 vectors as genetic tools for single copy gene cloning in *Acinetobacter baumannii*. J Microbiol Methods [Internet]. 2010;82(3):296–300. Available from: 10.1016/j.mimet.2010.07.002

5. Aranda J, Poza M, Pardo BG, Rumbo S, Rumbo C, Parreira JR, et al. A rapid and simple method for constructing stable mutants of *Acinetobacter baumannii*. BMC Microbiol [Internet]. 2010;10(1):279. Available from: http://www.biomedcentral.com/1471-2180/10/279

6. Stolze N, Bader C, Henning C, Mastin J, Holmes AE, Sutlief AL. Automated image analysis with ImageJ of yeast colony forming units from cannabis flowers. J Microbiol Methods [Internet]. 2019;164(July):105681. Available from: 10.1016/j.mimet.2019.105681

7. Geisinger E, Isberg RR. Antibiotic modulation of capsular exopolysaccharide and virulence in Acinetobacter baumannii. PLoS Pathog. 2015;11(2):1–27.

8. Geisinger E, Huo W, Hernandez-Bird J, Isberg RR. *Acinetobacter baumannii*.: envelope determinants that control drug resistance, virulence, and surface variability. Annu Rev Microbiol. 2019;73(1):481–506.

9. Niu T, Guo L, Luo Q, Zhou K, Yu W, Chen Y, et al. *wza* gene knockout decreases *Acinetobacter baumannii* virulence and affects Wzy-dependent capsular polysaccharide synthesis. Virulence [Internet]. 2020;11(1):1–13. Available from: 10.1080/21505594.2019.1700659

10. Tickner J, Hawas S, Totsika M, Kenyon JJ. The Wzi outer membrane protein mediates assembly of a tight capsular polysaccharide layer on the *Acinetobacter baumannii* cell surface. Sci Rep [Internet]. 2021;11(1):1–12. Available from: 10.1038/s41598-021-01206-5

11. Lees-Miller RG, Iwashkiw JA, Scott NE, Seper A, Vinogradov E, Schild S, et al. A common pathway for *O*-linked protein-glycosylation and synthesis of capsule in *Acinetobacter baumannii*. Mol Microbiol. 2013;89(5):816–30.

12. Valcek A, Nesporova K, Whiteway C, De Pooter T, De Coster W, Strazisar M, et al. Genomic Analysis of a Strain Collection Containing Multidrug-, Extensively Drug-, Pandrug-, and Carbapenem-Resistant Modern Clinical Isolates of Acinetobacter baumannii. Antimicrob Agents Chemother. 2022;66(9).

13. Singh JK, Adams FG, Brown MH. Diversity and function of capsular polysaccharide in *Acinetobacter baumannii*. Front Microbiol. 2019;10(JAN):1–8.

14. Hu L, Shi Y, Xu Q, Zhang L, He J, Jiang Y, et al. Capsule thickness, not biofilm formation, gives rise to mucoid *Acinetobacter baumannii* phenotypes that are more prevalent in long-term infections: A study of clinical isolates from a hospital in china. Infect Drug Resist. 2020;13:99–109.

15. Bai J, Dai Y, Farinha A, Tang AY, Syal S, Vargas-Cuebas G, et al. Essential Gene Analysis in *Acinetobacter baumannii* by High-Density Transposon Mutagenesis and CRISPR Interference. ASM -Journal Bacteriol. 2021;203(12):e00565–20.

16. Ellison CK. Bacterial adhesion at the single-cell level. Nat Rev Microbiol [Internet]. 2018;16(October). Available from: 10.1038/s41579-018-0057-5

17. Sauer K, Stoodley P, Goeres DM, Burmølle M, Stewart PS, Bjarnsholt T. The biofilm life cycle : expanding the conceptual model of biofilm formation. Nat Rev Microbiol. 2022;20(October).

18. Sahly H, Podschun R, Oelschlaeger TA, Greiwe M, Parolis H, Hasty D, et al. Capsule impedes adhesion to and invasion of epithelial cells by *Klebsiella pneumoniae*. Infect Immun. 2000;68(12):6744–9.

19. Béchon N, Mihajlovic J, Vendrell-Fernández S, Chain F, Langella P, Beloin C, et al. Capsular polysaccharide cross-regulation modulates *Bacteroides thetaiotaomicron* biofilm formation. MBio. 2020;11(3):1–14.

20. Spinosa MR, Progida C, Talà A, Cogli L, Alifano P, Bucci C. The *Neisseria meningitidis* capsule is important for intracellular survival in human cells. Infect Immun. 2007;75(7):3594–603.

21. Häuser S, Wegele C, Stump-Guthier C, Borkowski J, Weiss C, Rohde M, et al. Capsule and fimbriae modulate the invasion of *Haemophilus influenzae* in a human blood-cerebrospinal fluid barrier model. Int J Med Microbiol [Internet]. 2018;308(7):829–39. Available from: 10.1016/j.ijmm.2018.07.004

22. Davey ME, Duncan MJ. Enhanced biofilm formation and loss of capsule synthesis: Deletion of a putative glycosyltransferase in *Porphyromonas gingivalis*. J Bacteriol. 2006;188(15):5510–23.

23. Zhang W, Yao Y, Zhou H, He J, Wang J, Li L, et al. Interactions between host epithelial cells and *Acinetobacter baumannii* promote the emergence of highly antibiotic resistant and highly mucoid strains. Emerg Microbes Infect [Internet]. 2022;11(1):2556–69. Available from: 10.1080/22221751.2022.2136534

24. Tan YH, Chen Y, Chu WHW, Sham LT, Gan YH. Cell envelope defects of different capsule-null mutants in K1 hypervirulent *Klebsiella pneumoniae* can affect bacterial pathogenesis. Mol Microbiol. 2020;113(5):889–905.

25. Rendueles O. Deciphering the role of the capsule of *Klebsiella pneumoniae* during pathogenesis: A cautionary tale. Mol Microbiol. 2020;113(5):883–8.

26. Sanchez-Larrayoz AF, Elhosseiny NM, Chevrette MG, Fu Y, Giunta P, Spallanzani RG, et al. Complexity of Complement Resistance Factors Expressed by *Acinetobacter baumannii* Needed for Survival in Human Serum. J Immunol. 2017;199(8):2803–14.

27. Cahill SM, Hall RM, Kenyon JJ. An update to the database for *Acinetobacter baumannii* capsular polysaccharide locus typing extends the extensive and diverse repertoire of genes found at and outside the K locus. Microb Genomics. 2022;8(10):1–21.

28. Schembri MA, Dalsgaard D, Klemm P. Capsule Shields the Function of Short Bacterial Adhesins. J Bacteriol. 2004;186(5):1249–57.

29. Petruzzi B, Dickerman A, Lahmers K, Scarratt WK, Inzana TJ. Polymicrobial Biofilm Interaction Between *Histophilus somni* and *Pasteurella multocida*. Front Microbiol. 2020;11(July):1–14.

30. Maure A, Robino E, Henst C Van Der. The intracellular life of *Acinetobacter baumannii*. Trends Microbiol [Internet]. 2023;xx(xx):1–13. Available from: 10.1016/j.tim.2023.06.007

31. Rubio T, Gagné S, Debruyne C, Dias C, Cluzel C, Mongellaz D, et al. Incidence of an Intracellular Multiplication Niche among *Acinetobacter baumannii* Clinical Isolates. mSystems. 2022;7(1).

32. Sycz G, Venanzio G Di, Distel JS, Sartorio MG, Le NH, Scott NE, et al. Modern *Acinetobacter baumannii* clinical isolates replicate inside spacious vacuoles and egress from macrophages. PLoS Pathog [Internet]. 2021;17(8):1–19. Available from: 10.1371/journal.ppat.1009802

33. Wen Z, Zhang JR. Bacterial capsules [Internet]. Vols. 1–3, Molecular Medical Microbiology: Second Edition. Elsevier Ltd; 2014. 33–53 p. Available from: 10.1016/B978-0-12-397169-2.00003-2

34. Taylor C RI. The regulation of capsule expression. In: In: Wilson M, editor Bacterial Adhesion to Host Tissues: Mechanisms and Consequences. Cambridge: Cambridge University Press; 2002. p. 115–38.

35. Chin CY, Tipton KA, Farokhyfar M, Burd EM, Weiss DS, Rather PN. A high-frequency phenotypic switch links bacterial virulence and environmental survival in *Acinetobacter baumannii*. Nat Microbiol [Internet]. 2018;3(5):563–9. Available from: 10.1038/s41564-018-0151-5

